# Full-Length Molecular Models of Brain-Derived α-Synuclein Fibrils Reveal a Fuzzy-Coat-Mediated Mechanism for Selective Peptide Binding

**DOI:** 10.64898/2026.04.16.718707

**Authors:** Carlos Pintado-Grima, Oriol Bárcenas, Giulio Tesei, F. Emil Thomasen, Kresten Lindorff-Larsen, Salvador Ventura

## Abstract

Parkinson’s disease (PD) is characterized by the aggregation of α-synuclein (aSyn) into amyloid fibrils that seed further aggregation and contribute to pathological spreading. Peptides that bind aggregated aSyn are promising therapeutic leads, but their validation is slow, difficult to standardize, and often relies on structural models limited to the ordered cross-β core. Here, we built models of brain-derived full-length aSyn fibrils by extending the Lewy-fold cryo-EM structure with disordered N- and C-terminal segments and sampling the resulting ensembles with the CALVADOS coarse-grained force field. The resulting fibrils display a dynamic fuzzy coat in which the termini, especially the acidic C-terminal tails, form recurrent transient contacts with the core, including the aggregation-prone β5 and β9 motifs. We then used these full-length fibrils in a standardized *in silico* assay for peptide binding. Simulations of the validated peptide binders PSMα3 and LL-37 reproduced their relative binding behavior and converged on a common mechanism in which electrostatic capture by the anionic fuzzy coat precedes stabilization on recurrent P2 and P3 hotspots within the structured core. Control simulations with monomeric aSyn or core-only fibrils showed that persistent association is lost in the absence of the full-length architecture, providing a mechanism for selectivity toward aggregated species. Finally, screening 123 peptides from aSynPEP-DB using a relative contact-based binding score yielded a ranked set of candidate binders and identified net positive charge as the dominant determinant of sustained association, with hydrophobicity acting as a secondary modulator. Together, these results establish full-length, brain-derived fibril ensembles as a practical framework for understanding ligand recognition at pathological amyloid surfaces and for prioritizing therapeutic peptide binders targeting aggregated aSyn.

**Significance:** Parkinson’s disease is driven by the assembly of α-synuclein into amyloid fibrils, yet most structural models of these aggregates omit the disordered termini that form the fibrillar “fuzzy coat” *in vivo*. Here we use coarse-grained simulations to reconstruct full-length, brain-derived α-synuclein fibrils and show that this fuzzy coat transiently contacts the Lewy-fold core, reshaping access to cross-β surface motifs. Using these ensembles in a computational assay, we recapitulate the relative binding behavior of validated peptide inhibitors and reveal a two-step mechanism in which cationic and amphipathic peptides are first captured by the anionic fuzzy coat and then engage recurrent core hotspots. This framework explains selective recognition of aggregated α-synuclein and provides a practical route to prioritize therapeutic peptide binders.

## Introduction

Parkinson’s disease (PD) is the fastest-growing neurodegenerative disorder worldwide (1, 2). Although dopaminergic treatments provide meaningful clinical symptomatic benefit, they do not halt the progressive neurodegeneration that characterizes the disease (3). Developing disease-modifying strategies, therefore, requires targeting the molecular processes that drive PD pathogenesis.

A defining pathological hallmark of PD and related synucleinopathies is the aggregation of α-synuclein (aSyn) into toxic oligomers and amyloid fibrils (4–6). In solution, aSyn is an intrinsically disordered protein (7) composed of an amphipathic N-terminal region, a hydrophobic non-amyloid-β component (NAC) segment that nucleates amyloid formation, and a highly acidic C-terminal tail (8). During aggregation, aSyn populates a heterogeneous continuum of assemblies that differ in size, structure, and toxicity (9, 10). Among these species, amyloid fibrils accumulate prominently in aged brains and serve as catalytic surfaces for secondary nucleation, accelerating fibril amplification and facilitating pathological spreading within neural tissue (11).

Amyloid fibrils share a characteristic cross-β molecular architecture, in which β-strands stack perpendicular to the fibril axis to form elongated protofilaments that may associate laterally (12, 13). Recent cryo-electron microscopy studies have resolved atomic structures of several aSyn fibril polymorphs, including fibrils extracted directly from patient brain tissue (14, 15). However, even high-resolution reconstructions typically resolve only the ordered cross-β core. The N- and C-terminal regions remain invisible because they are conformationally heterogeneous in the fibrillar state. *In vivo*, these unresolved segments form a dynamic “fuzzy coat” surrounding the amyloid core (16).

For aSyn fibrils, this fuzzy coat is expected to be especially prominent. Nearly half of the sequence remains disordered in the fibrillar state, including the 40-residue acidic C-terminal tail that generates a highly anionic surface around the fibril. Consequently, the effective molecular interface presented by pathological fibrils is defined not only by the ordered cross-β scaffold but also by fluctuating disordered segments (17) that shape accessibility, electrostatics, and intermolecular recognition. Evidence from other amyloid systems supports the functional importance of these disordered layers. Studies of Aβ42 and Tau fibrils have shown that fuzzy coats can establish recurrent contacts with the structured core, thereby modulating solvent accessibility and local solubility (18, 19). These observations suggest that the dynamic outer layer of amyloid fibrils may strongly influence how molecular ligands, chaperones (20) or receptors (21) recognize aggregated species.

This issue is particularly relevant for therapeutic ligand discovery. Structural models that omit the fuzzy coat are likely to misrepresent the binding landscape encountered by ligands targeting aggregated proteins. Peptides represent attractive modulators of amyloid assembly because they can engage extended protein surfaces with high specificity and can be rationally tuned through sequence-level modifications (22). In the case of aSyn, we previously identified short peptides that bind toxic oligomers and fibrils with low-nanomolar affinity while showing negligible recognition of the functional monomeric state, including the bacterial peptide PSMα3 and the human host-defense peptide LL-37 (23, 24). These observations indicate that aggregation generates a physicochemically distinct binding surface that can be selectively recognized by peptides. However, the molecular determinants underlying this selectivity remain poorly understood, and the potential role of the fuzzy coat in shaping ligand accessibility to the fibril surface has not been addressed.

At the same time, experimental validation of peptide binders remains slow and difficult to standardize. This limitation is increasingly important as the number of candidate aSyn-binding peptides continues to expand. For example, the aSynPEP-DB database catalogs more than one hundred peptide sequences reported to interact with aggregated aSyn (25), several of which have shown evidence of alleviating PD-related phenotypes in cellular or animal models (26). A consistent *in silico* framework based on complete fibril structures could therefore complement experimental screening and help rationalize the mechanisms of candidate ligands.

Here we place full-length, brain-derived aSyn fibrils at the center of peptide-binder validation. Starting from a patient-derived Lewy-fold core structure (15), we reconstruct full-length fibrils and characterize the resulting fuzzy coat and its transient interactions with the structured cross-β scaffold using simulations with the CALVADOS coarse-grained model. This implicit-solvent framework has been parameterized using experimental data for intrinsically disordered and multidomain proteins and has proven effective in describing the conformational behavior of disordered protein assemblies (27, 28) and proteins containing mixtures of ordered and disordered regions (29, 30). By restoring the sequence regions absent from cryo-EM structures, these models allow us to examine how disordered terminal segments shape the physicochemical environment surrounding the fibril core.

Using ensembles of these full-length fibrils as a standardized simulation platform, we interrogate peptide binding to aggregated aSyn. Our results reveal that the fuzzy coat modulates accessibility of aggregation-prone surface motifs and shapes the electrostatic landscape encountered by approaching ligands. The simulations reproduce the relative binding trends observed experimentally for previously validated peptides and uncover a two-step binding mechanism in which peptide capture by the disordered fuzzy coat precedes engagement of hotspots on the fibril core. Together, these findings establish full-length, brain-derived fibril models as a practical framework for understanding amyloid-fibril surface chemistry and for mechanistically validating peptide ligands targeting aggregated aSyn.

## Results

### Structures of brain-derived aSyn fibrils and full-length fibril modeling

To work with a disease-relevant scaffold, we selected the cryo-EM structure of α-synuclein (aSyn) filaments purified from brains of patients with Parkinson’s disease (PD), Parkinson’s disease dementia (PDD), and dementia with Lewy bodies (DLB) (15). These filaments were reported to be structurally indistinguishable across the three conditions and to consist of a single protofilament defining the “Lewy fold” polymorph (PDB: 8A9L). The ordered cross-β core spans residues 31–100 and forms nine β-strands (β1–β9) arranged in a three-layered architecture (β1–β5, β6–β8, and β9). Using the helical rise and twist reported for this polymorph (4.76 Å and 0.86° per subunit, respectively), we propagated the protofilament to generate a fibril segment of 209 stacked monomers (≈100 nm along the fibril axis), corresponding to ∼180° cumulative rotation and therefore capturing half a helical turn **(Fig. 1A).** This length provides a computationally tractable fibril segment for coarse-grained simulations while reducing finite-size artifacts associated with fibril ends.

**Fig. 1.**
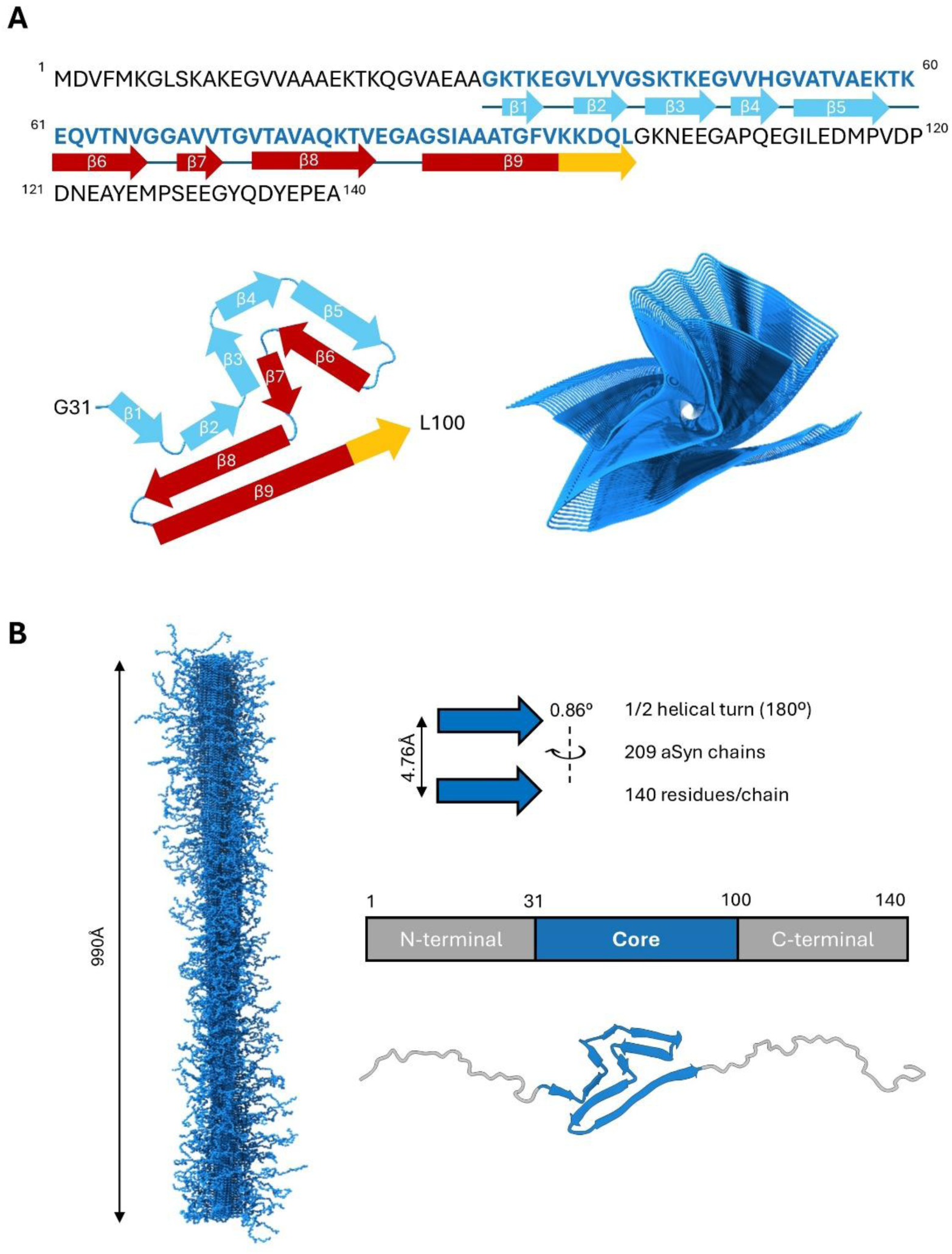
Construction of full-length, brain-derived aSyn fibril models. (A) aSyn sequence and top views of the cryo-EM structure of the PD/DLB aSyn protofilament core (residues 31–100; PDB: 8A9L). The long fibril was reconstructed using the reported helical parameters, with a reconstructed core fibril showing a 180° twist. Blue, red and ochre colors of β-strands indicate overlap with N-terminal, NAC and C-terminal regions. (B) Full-length fibril model after building N- and C-terminal disordered segments for each chain, yielding a fuzzy coat surrounding the cross-β core.

The atomic model of the Lewy fold resolves only the structured core; while the N- and C-terminal segments are present in the fibrils, they are not visible in the cryo-EM density, consistent with their intrinsic disorder and conformational heterogeneity in the fibrillar state. These flanking regions form a dynamic “fuzzy coat” that can modulate fibril surface chemistry, steric accessibility, and make transient contacts with solvent-exposed regions of the cross-β core (18, 31). To obtain a full-length fibril model, we appended residues 1–30 and 101–140 to each chain with MODELLER, generating a cross-β core surrounded by disordered termini. We then relaxed and sampled the conformational ensemble of these termini using molecular dynamics simulations with the coarse-grained CALVADOS model while maintaining the experimental Lewy-fold geometry through elastic network restraints on the core residues. The resulting conformational ensemble captures a dynamic fuzzy coat extending into solution (**Fig. 1B**), providing a physically plausible representation of the surface presented by brain-derived aSyn fibrils *in vivo*.

### The fuzzy coat establishes contact with the fibrillar core

The fuzzy coat created by the N- and C-terminal disordered regions holds the potential to contact the cross-β core. In simulations of the full-length fibril, both termini formed transient but recurrent contacts with the core. Contact map analysis reveals the identities of these interactions (**Figs. 2A and 2B and SI Fig. S1**). First, the N-terminal disordered region engages the C-terminal portion of the ordered core, specifically the NAC hinge between β8 and β9. Second, the acidic C-terminal tail forms contacts with multiple solvent-accessible core elements, including β3–β5 and β9. Overall, C-terminal interactions dominate the fuzzy-coat/core contact network, accounting for three-quarters (73.2%) of all tail-to-core contacts (**Fig. 2C**)

**Fig. 2.**
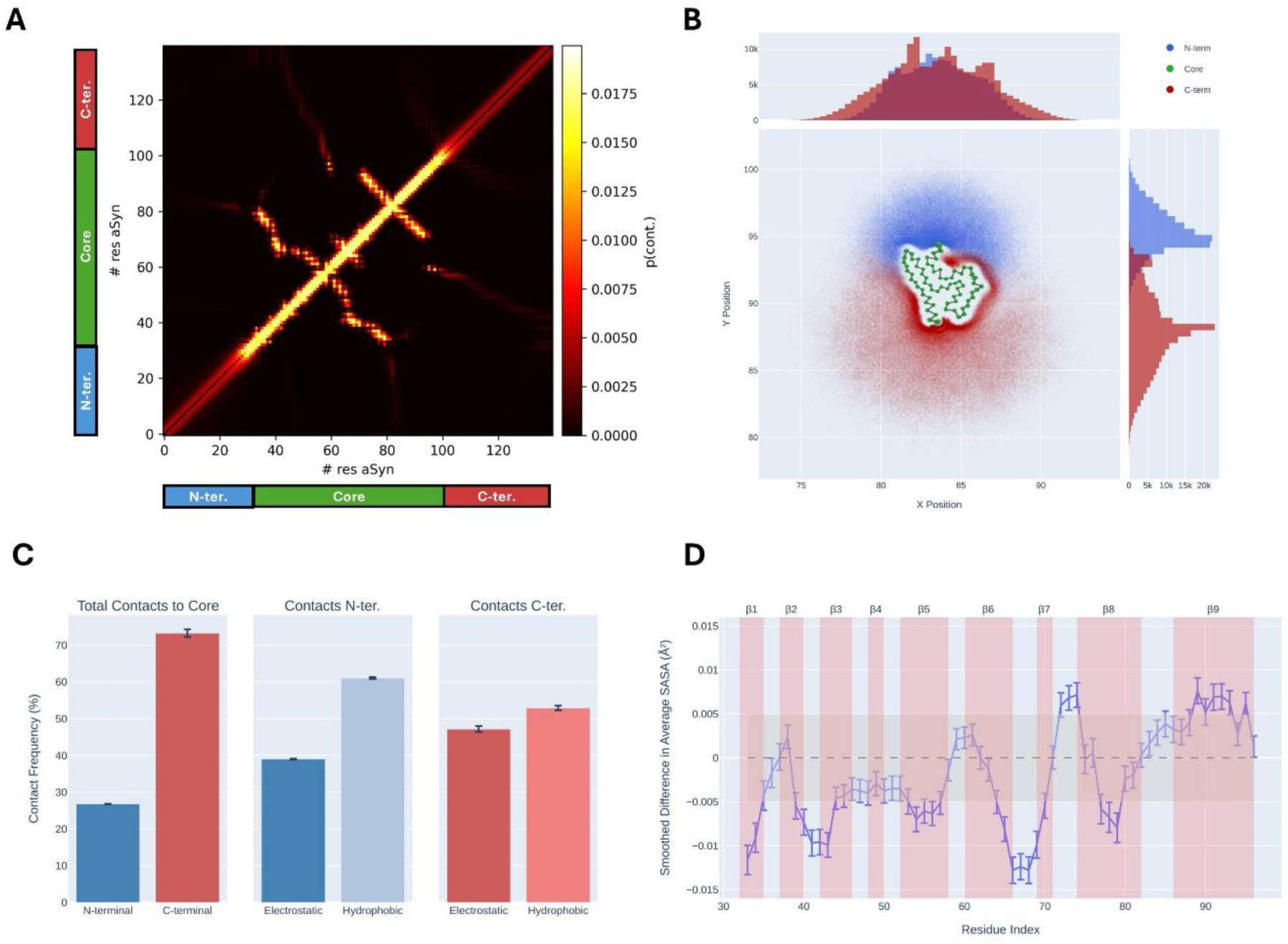
Transient fuzzy coat–core contacts shape the fibril surface. (A) Intrafibril contact map from fibril-only simulations highlighting transient contacts between disordered termini and the structured core. (B) Average spatial positions of aSyn residues over the trajectory colored by N-terminal, core and C-terminal regions (blue, green and red, respectively). (C) Contact frequency and contact types for both N- and C- terminal IDRs with the cross-β core. (D) Per-residue SASA for the fibril core region, computed from the central portion of the fibril to avoid artifactually high solvent exposure from chains in the extremes; the fuzzy coat reduces solvent exposure of specific residues relative to a core-only model. The grey background indicates a ±0.005 margin of error as a confidence interval. The light-red background indicates the location of β-strands. Error bars for panels (C) and (D) were computed as the standard error of the mean (SEM) obtained by grouping all chains in the amyloid fibril into segments of 8 chains.

N-terminal/core contacts are dominated by hydrophobic interactions (61.0%), whereas C-terminal contacts show a more balanced contribution from electrostatic (47.1%) and hydrophobic (52.9%) interactions (**Fig. 2C**). These intrafibril contacts were not static; rather, different chains and time windows contributed, consistent with a rapidly interconverting ensemble of disordered conformations. In this way, the dynamic nature of the C-terminal IDR covers and protects most of the first and third β-strand layers, which are the only layers accessible to solvent in the core, since the second layer remains essentially buried by the Lewy fold architecture.

We next performed a solvent-accessible surface area (SASA) analysis. We computed the SASA per residue of the protofilament cross-β in the absence and presence of the fuzzy coat (**Fig. 2D**). We observed a more marked reduction of the SASA for core residues mapping to β1, β2-β3, β5, β6-β7 and β8, consistent with partial shielding of the core surfaces by disordered termini. The changes were modest on an absolute scale, supporting a “dynamic shielding” mechanism in which the fuzzy coat intermittently occupies core epitopes rather than permanently burying them.

### Overlap of fibril core aggregation-prone motifs and fuzzy-coat contact regions

We then examined if the identified contacts overlap with sequence-encoded aggregation-prone regions (APRs) in the cross-β core. AGGRESCAN (32) analysis of the full-length aSyn sequence showed a enrichment in aggregation propensity within the structured core region (residues 31–100), with four major APRs located at β2, β5, the region between the end of β6 and the start of β8, and the long terminal β9 strand. In contrast, the fuzzy coat sequence regions exhibited much lower aggregation propensity (**Fig. 3A**). This aligns with our recent findings that sequence-based predictors effectively identify the regions that most often form the structural core of aSyn polymorphs (33).

**Fig. 3.**
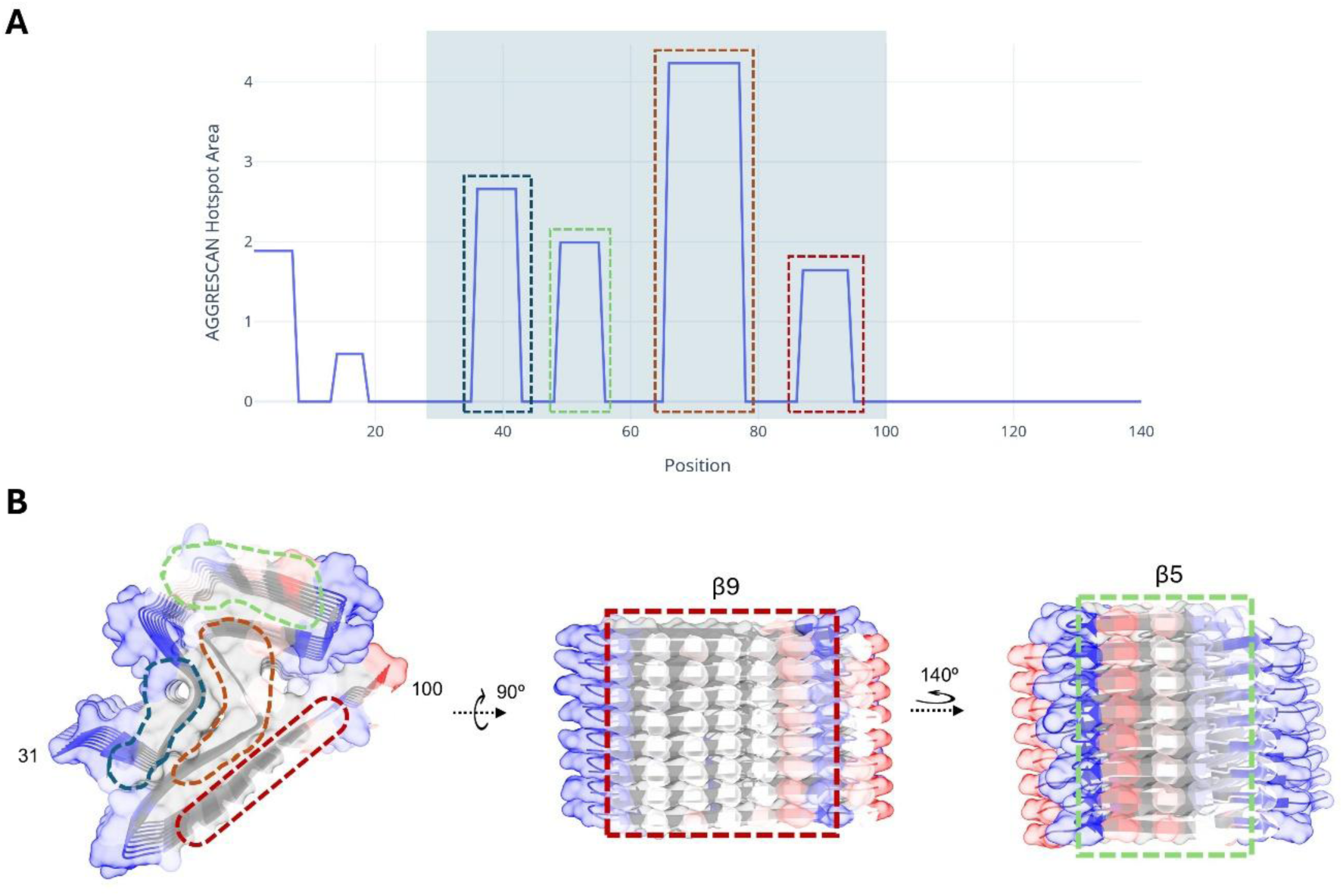
Aggregation-prone motifs define interaction-prone core patches. (A) Aggregation hotspots predicted by consecutive aggregation-prone segments by AGGRESCAN. The grey background highlights the structured core region. (B) Top and lateral views of the structural aggregation propensity predicted by Aggrescan4D for eight core chains extracted from the fibril center; red indicates aggregation-prone regions, blue indicates more soluble regions and grey indicates no solvent exposition.

We then mapped these sequence hotspots onto their structural context by computing the structural aggregation propensity (STAP) of the Lewy-fold core using Aggrescan4D (34). In contrast with the sequence-based prediction by AGGRESCAN, Aggrescan4D allows for the identification of STAPs on the surface of globular domains. This analysis highlights a key separation between intrinsic aggregation propensity and effective aggregation risk on the fibril surface (**Fig. 3B**): APRs located in β2 and in the β6–β8 region are largely buried within the interior of the three-layered core, whereas β5 and β9 form two solvent-exposed hydrophobic “ladders” on opposite faces of the protofilament. Notably, these two exposed patches coincide with the dominant sites contacted by the C-terminal tails in the full-length simulations. It is worth noting that, in our calculations, the STAP of these ladders is moderate. In addition, the connecting regions between β-strands are predicted to be significantly soluble (**Fig. 3B**). Together, this makes the overall STAP of the fibril core surface low compared to that of other pathogenic fibrils, such as Aβ42 (**SI Fig. S2**). This might explain why, when we used Aggrescan4D to recalculate the STAP of the fibrillar core in the presence of the fuzzy coat, despite the observed contacts, the core residues did not display decreased aggregation scores (**SI Fig. S3 and SI Table S1**), as reported for Aβ42 (18). Instead, the primary effect is a local reduction in predicted solubility for the polar β-sheet connecting segments, which become partially shielded by the fluctuating termini, mirroring the calculated SASA reductions for these regions.

### Full-length fibrils as a standardized *in silico* assay reproduce binding trends of validated peptides

We recently identified a family of short peptides that bind aSyn fibrils with affinity in the low nanomolar range and clear conformational selectivity, showing negligible recognition of the functional, intrinsically disordered monomer (23). This selectivity indicates that the relevant binding determinants emerge only upon assembly, where fibril formation reorganizes the physicochemical landscape of aSyn. Simulations suggest that assembly generates persistent hydrophobic ladders embedded within a highly anionic environment, largely shaped by the disordered C-terminal tail, which carries a formal net charge of −12 and concentrates 15 Asp/Glu residues. This architecture produces a diffuse yet physicochemically defined binding surface, expected to favor ligands that combine electrostatic complementarity (to promote capture/retention within the polyelectrolyte brush) with hydrophobic patterning (to stabilize contacts on exposed core patches). Consistent with this model, we showed that PSMα3, a 22-residue cationic, amphipathic, bacterial peptide, inhibits aSyn aggregation and binds fibrils with a *K*_d_ of 7.8 nM while showing no detectable interaction with monomeric aSyn. Indeed, a fluorescently labeled derivative of PSMα3 was subsequently shown to detect aSyn aggregates *in vivo* with high accuracy, selectively labeling aggregated over monomeric species across gastrointestinal and brain tissues (35).

To test whether simulations based on our brain-derived fibril models can be used to recapitulate binding trends and provide a molecular description of ligand recognition, we used CALVADOS simulations of full-length Lewy-fold fibrils with the addition of free peptides in solutions as a standardized *in silico* assay for peptide binding to the fibril. We benchmarked the assay with PSMα3 and simulated a 2:1 ratio of fibril subunits to peptide molecules to account for the fact that the peptide totally abrogates fibril formation at this ratio (23).

In the production simulation (SP-PSM), PSMα3 formed persistent contacts with the fibril surface. Contact maps revealed two dominant interaction features: strong contacts with the acidic C-terminal tails and two recurrent patches within the fibril-structured core (**Fig. 4A**). The first core interaction site spans residues 49–57 and maps to β5, overlapping with the previously described P2 region, which acts as a “master controller” of aSyn aggregation. We showed that P2 deletion strongly inhibits amyloid formation by impeding the oligomer-to-fibril transition (36). The second patch spans residues 84–93 and maps to β9. Because this surface element repeatedly emerged as an interaction site for both the fuzzy coat and peptide ligands in our simulations, we refer to it here as P3. As indicated above, P2 and P3 form the only solvent-exposed aggregation-prone ladders on opposite faces of the fibril core. The spatial coincidence between fuzzy-coat contacts and peptide binding sites suggests that the same surface regions that dynamically interact with the disordered tails remain accessible to external ligands. Across trajectories, multiple PSMα3 molecules associated intermittently with these fibril sites, indicating a multivalent and thermodynamically favorable binding landscape (**Fig. 4B**).

**Fig. 4.**
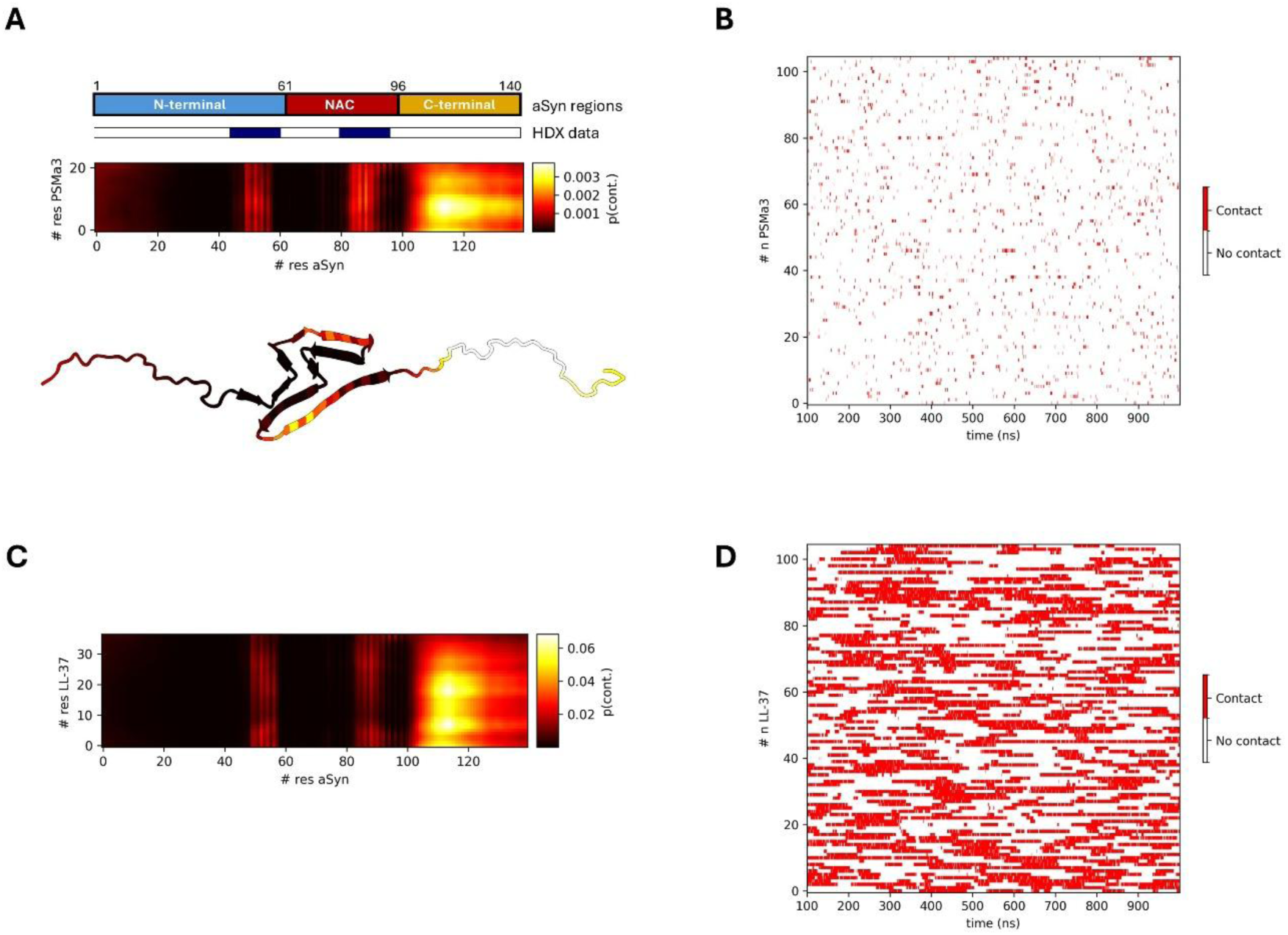
Full-length fibril simulations reproduce binding patterns of validated peptide inhibitors. (A) Contact map between PSMα3 and the full-length fibril showing interactions with the C-terminal tails and two recurrent core patches (P2 and P3). The color of the fibril representation indicates the per-residue probability of contact with PSMα3, using the same palette as the contact map. (B) Time series of the number of PSMα3 peptides bound. (C) Contact map between LL-37 and the full-length fibril showing the same recurrent sites. (D) Time series of the number of LL-37 peptides bound.

Direct mapping of PSMα3 epitopes on mature fibrils by HDX-MS was not feasible in our previous work because fibrils are highly resistant to proteolysis, but we mapped the interaction on aSyn oligomers to which the peptide binds with a *K*_d_ of 6.7 nM, closely matching its affinity for fibrils. Interestingly, the only regions showing reduced deuterium uptake corresponded to the P2 and P3 segments. Moreover, MAS-ssNMR measurements of ^13^C^15^N labelled aSyn oligomers indicated that the rigid core residues 70–89 in the oligomer occupy a chemical environment similar to that in fibrils, with chemical shift differences < 1 ppm, supporting continuity of key structural elements between these assemblies. Together, these observations suggest that PSMα3 engages conserved core-adjacent APRs that are shared by oligomers and fibrils.

To further evaluate whether the simulation assay captures the physicochemical logic established experimentally, rather than overfitting to a specific sequence, we simulated two PSMα3 variants previously used to dissect determinants of inhibition (23). A variant in which all hydrophobic residues were substituted by Leu retained inhibitory activity close to wild type experimentally; in our simulations, this Leu-substituted peptide retained the same dominant interaction pattern and contacted the same fibril regions as wild type (**SI Fig. S4A and S4B**). In contrast, a charge-reversal variant (K6E/K12E), which shifts net charge from cationic to anionic and abolishes interaction/inhibition experimentally, showed no stable association with the fibril surface under identical simulation conditions (**SI Fig. S4C**). Together, these results support a model in which net positive charge is required for capture by the anionic fuzzy coat, while amphipathic patterning enables stabilization on exposed core ladders once proximity is achieved.

We then simulated the interactions of LL-37, a human antimicrobial peptide with similar amphipathic and cationic features and reported inhibitory activity against aSyn aggregation (24). In the production simulation (SP-LL37), LL37 showed similar interaction patterns to PSMα3: dominant contacts with C-terminal tails accompanied by repeated engagement of the P2 and P3 core patches (**Fig. 4C**). Relative to PSMα3, LL-37 displayed a 20-fold higher probability of association with the fibril (**Fig. 4D**), which is qualitatively consistent with its experimentally reported stronger affinity for both oligomers (*K*_d_= 3.62 nM) and fibrils (*K*_d_= 5.14 nM), respectively (23).

Overall, these results indicate that full-length fibril simulations with CALVADOS can capture relative binding trends and, importantly, map interactions onto physically realistic fibril surfaces.

### A two-step binding mechanism: electrostatic capture by the fuzzy coat followed by hotspot engagement on the cross-β core

To understand how peptides access structurally encoded hotspots within the full-length fibril architecture, we analyzed LL-37 binding dynamics in more detail. LL-37 contacts were broadly distributed along the fibril length, except near the fibril ends, where the reduced density of disordered tails lowers the local negative charge and diminishes electrostatic retention of cationic ligands (**Fig. 5A**).

**Fig. 5.**
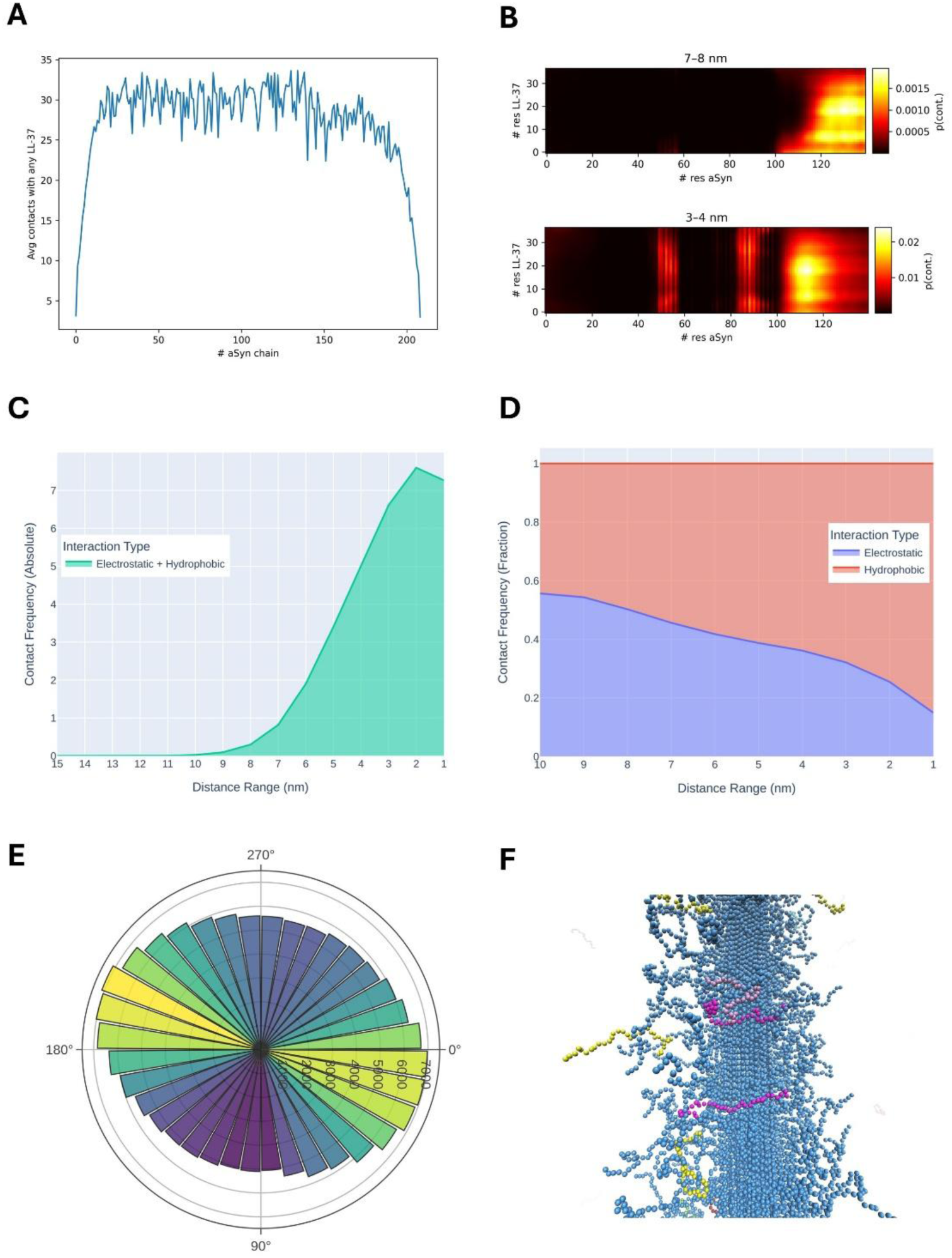
LL-37 binding follows a two-step mechanism: fuzzy coat capture and core hotspot engagement. (A) LL-37 contacts with individual aSyn chains across the fibril length. (B) Average LL-37–aSyn contact maps binned by peptide–fibril COM distance (7–8 nm versus 3–4 nm). (C) Absolute contact frequencies at different peptide-fibril distances. (D) Distance-dependent decomposition of contacts into electrostatic and hydrophobic interactions. (E) Preferred binding orientation of LL-37 relative to the fibril axis in the bound state. Green and yellow colors indicate higher binding probabilities. (F) Representative frame shot depicting the interaction of LL-37 peptides with the fuzzy coat (yellow peptides) and the core (pink peptides).

Distance-resolved contact analysis revealed a clear, sequential binding landscape (**Fig. 5B and SI Fig. S5**). At intermediate peptide–fibril distances (7–8 nm), LL-37 interacted predominantly with the outermost portion of the disordered C-terminal tails, yielding contact maps dominated by acidic tail residues. Upon closer approach (3–4 nm), LL-37 established frequent contacts with the P2 and P3 patches on the cross-β core while maintaining interactions with the inner segments of the C-terminal tails. Thus, the fuzzy coat does not simply act as a passive shield: it functions as an anionic capture layer that concentrates the peptide near the fibril surface and enables repeated attempts to engage exposed hydrophobic core motifs. Notably, because the C-terminal tails themselves transiently sample the P2/P3 surfaces in the full-length ensemble, tail–core dynamics provide a natural route by which a peptide captured by the fuzzy coat can be brought into proximity of these core hotspots.

The distance-dependent decomposition of contacts revealed first interactions happening at <10 nm (**Fig. 5C**) and a corresponding shift in interaction type (**Fig. 5D**). At larger distances, contacts were dominated by electrostatic interactions between basic residues in LL-37 and acidic residues in the fuzzy coat, given the fuzzy coat net charge (NC) per aSyn molecule of -12 (exceeding -900 in our modelled protofibril segment) and the NC of each peptide being +6. Once LL-37 approached the core, hydrophobic contacts increased in relative frequency, indicating engagement with APRs in the core. Together, these trends support a mechanism in which electrostatic capture and confinement by the fuzzy coat precede, and facilitate, mixed electrostatic/hydrophobic stabilization on core hotspots.

Finally, we asked whether productive binding is accompanied by a preferred peptide orientation on the anisotropic cross-β surface. In the bound regime (<5 nm), LL-37 preferentially aligned approximately perpendicular to the fibril axis and roughly parallel to the aSyn chain direction (**Figs 5E and 5F**), whereas at larger distances, the angular distribution was broad and approached random (**SI Fig. S6**). This emergence of orientational order upon binding is consistent with LL-37 engaging an extended, directionally structured surface rather than a localized pocket. Collectively, these analyses support a model in which peptides are first concentrated near the fibril by the anionic fuzzy coat, then penetrate inward and adopt a preferred bound geometry while engaging the solvent-accessible P2/P3 motifs on the structured core.

### Coupled conformational responses of the peptide and fuzzy coat upon binding

Although CALVADOS is a coarse-grained model, it is well-suited to quantify ensemble-level dimensions of disordered chains both alone and in context of structured regions (29). We therefore quantified the radius of gyration (*R*_g_) of aSyn C-terminal tails and LL-37 in the presence and absence of the cognate binding partner. In fibril-only simulations, the C-terminal tails populated relatively expanded conformations, as expected for a densely grafted acidic brush in which electrostatic repulsion favors extension into solution. Upon LL-37 binding, the tails compacted modestly (**Fig. 6A**), which is consistent with transient multivalent association of the cationic peptide with the acidic tails, which locally reduces tail–tail repulsion and promotes a more compact brush. Conversely, LL-37 peptides became more extended in the presence of the fibril (**Fig. 6B**), consistent with the idea that binding favors elongated peptide conformations. These coupled changes suggest a “mutual adaptation” between ligand and fuzzy coat, where peptide binding reorganizes the anionic brush, and the fuzzy coat selects peptide conformations that are more favorable for productive core engagement.

**Fig. 6.**
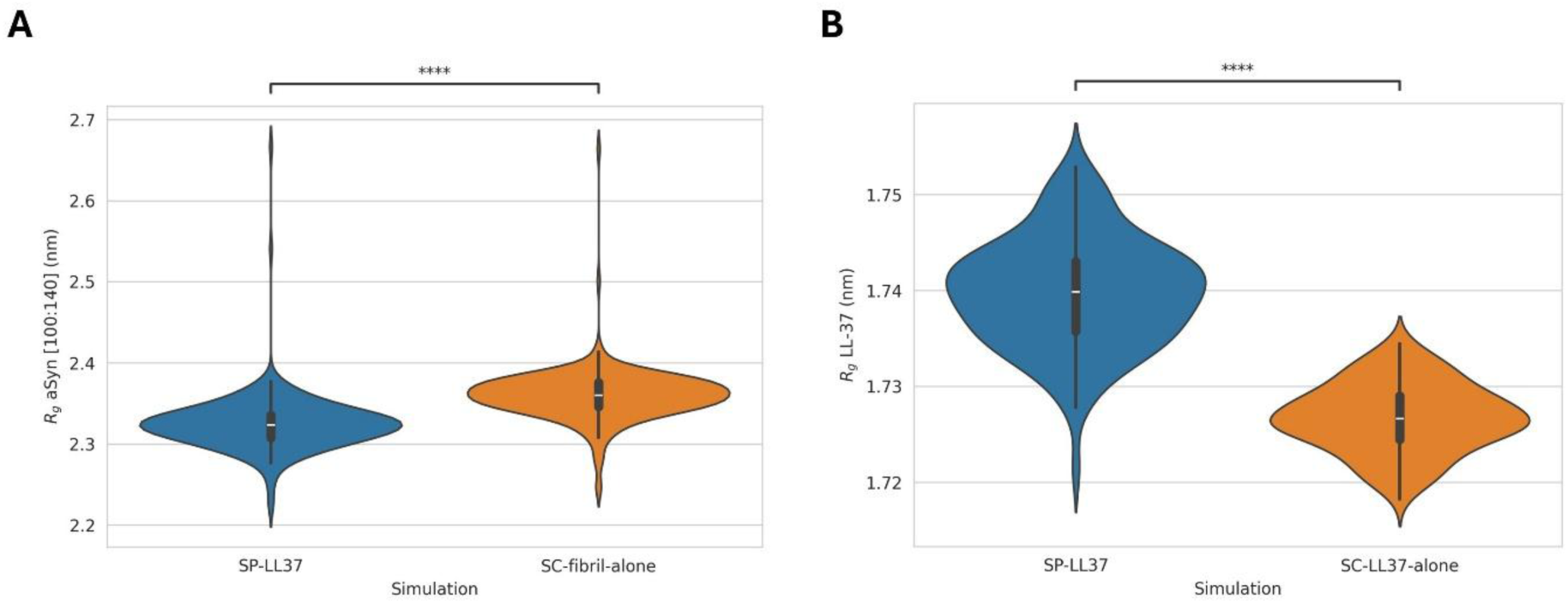
Coupled conformational responses of peptide and fuzzy coat. (A) Radius of gyration (*R*_g_) of aSyn C-terminal tails (residues 100–140) in the simulations in the presence (SP-LL37) and absence (SC-fibril-alone) of LL-37. (B) *R*_g_ of LL-37 peptides in the simulations in the presence (SP-LL37) and absence (SC-LL37-alone) of the full-length fibril. Nominally, **** indicates p<1e-32 for a difference in average *R*_g_. The simulated systems are described in Table 1.

**Table 1.** Summary and nomenclature of simulated systems.

| System name | Simulation type | Peptide entity | aSyn conformation |
| --- | --- | --- | --- |
| SP-PSM | Production | PSMα3 | Full-length fibril |
| SP-LL37 | Production | LL-37 | Full-length fibril |
| SC-all-Leu | Control | PSMα3 all-Leu variant | Full-length fibril |
| SC-anionic | Control | PSMα3 anionic variant | Full-length fibril |
| SC-monomer | Control | LL-37 | Monomeric aSyn (full-length, disordered) |
| SC-core | Control | LL-37 | Fibril core only (residues 31–100) |
| SC-helical | Control | LL-37 (helical) | Full-length fibril |
| SC-fibril-alone | Control | – | Full-length fibril |
| SC-LL37-alone | Control | LL-37 | – |
| SALL-pos | Screen | aSynPEP-DB peptides | Full-length fibril |
| SALL-neg-hel | Screen | Low-helicity negatives | Full-length fibril |
| SALL-neg-uh | Screen | Non-amphipathic negatives | Full-length fibril |
| SALL-neg-ncpr | Screen | Negatively charged negatives | Full-length fibril |

### Persistent binding requires the full-length fibril architecture

We next tested whether the full-length fibril architecture is necessary to reproduce persistent binding and selectivity. In simulations of LL-37 with monomeric aSyn treated as full-length disordered chains (SC-monomer), peptide encounters were infrequent and short-lived in comparison with the full-length fibril (**Figs. 7A and 7B**). Contacts occurred primarily with the acidic C-terminal segment but did not persist, consistent with the absence of a stable anionic brush and with the high entropic costs of maintaining multivalent contacts on a flexible monomer ensemble.

**Fig. 7.**
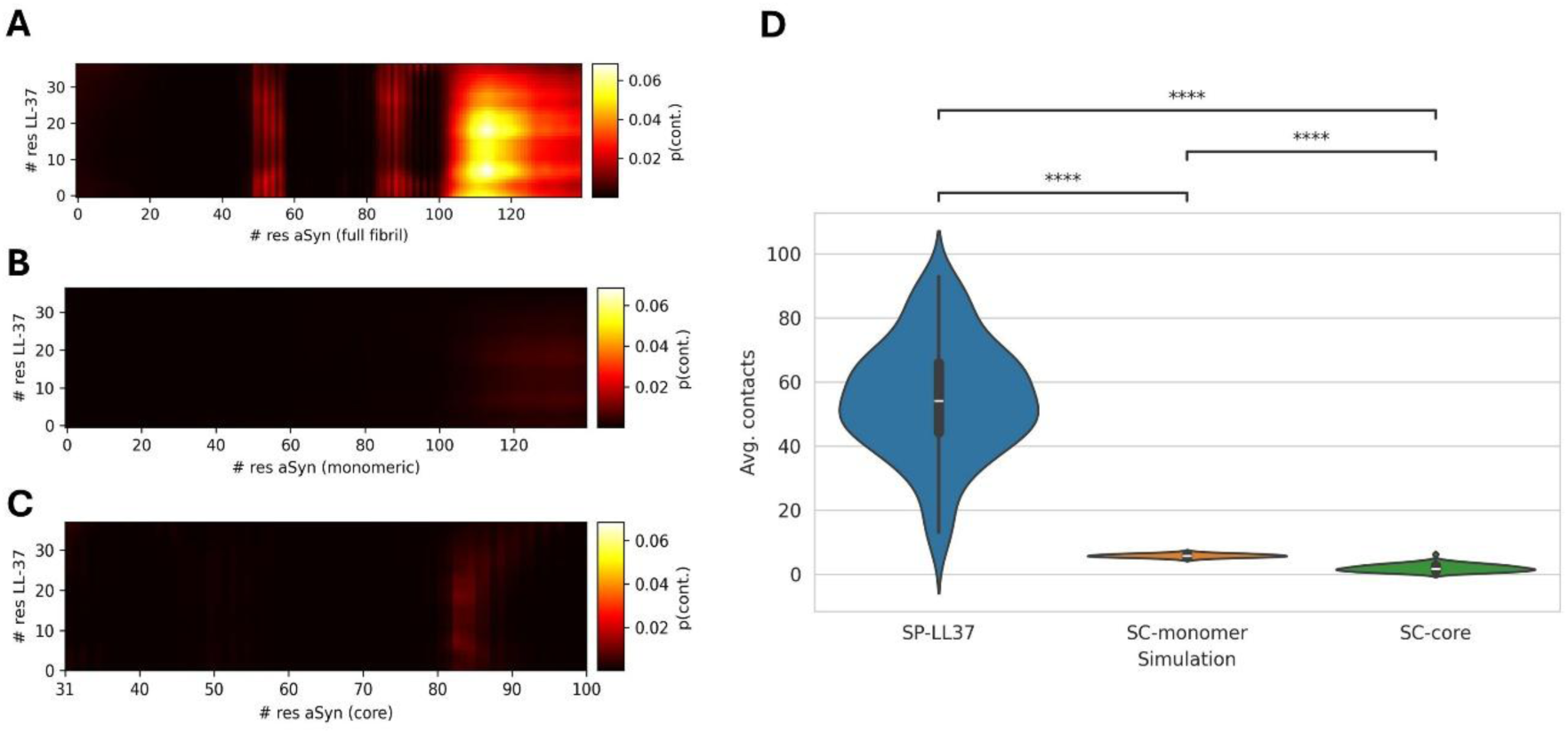
Persistent binding requires the full-length fibril architecture. (A) Contact map between LL-37 and full-length aSyn fibril (SP-LL37 simulation). (B) Contact map between LL-37 and monomer (SC-monomer simulation). (C) Contact map between LL37 and a core-only fibril lacking the fuzzy coat (SC-core simulation). (D) Average total contacts for the production simulation compared with control simulations; nominally, **** indicates p<5.454e-35 for a difference in average contacts. The simulated systems are described in Table 1.

We then isolated the contribution of the fuzzy coat by simulating LL-37 with a core-only fibril lacking the disordered termini (SC-core). In this system, LL-37 could still sample the structured cross-β surface and form occasional contacts with exposed hydrophobic elements, but these interactions were weak, transient, and largely restricted to P3/β9 (**Fig. 7C**). By contrast, productive engagement of P2/β5 was essentially lost. These results indicate that the fuzzy coat is not merely an additional binding surface appended to the core; rather, it provides an electrostatic capture and retention mechanism that keeps peptides in the vicinity of the core long enough to engage hotspot regions, especially for P2, whose efficient access depends on the full-length architecture (**Fig. 7D**).

### Full-length fibrils enable moderate throughput ranking of candidate binders and identify net positive charge as the dominant driver

To date, only PSMɑ3 and LL-37 have been experimentally validated to bind aSyn fibrils. However, a set of 123 additional candidate peptides with similar predicted properties has been compiled in aSynPEP-DB and awaits validation (25). Having established that full-length fibril simulations reproduce validated binding trends and yield a coherent mechanism of action, we applied the same assay to the candidate peptides in aSynPEP-DB. In a single simulation containing one copy of each of 123 peptides (SALL-pos), we quantified the average number of peptide–fibril contacts. We interpret this metric as a relative binding score for prioritization and ranked peptides accordingly (**Fig. 8A**). Notably, 79 of the 123 peptides (63.41%) formed more contacts than the benchmark binder PSMα3, and 14 exceeded LL-37, indicating that the database contains multiple peptides with strongly predicted propensity to associate with the full-length fibril surface.

**Fig. 8.**
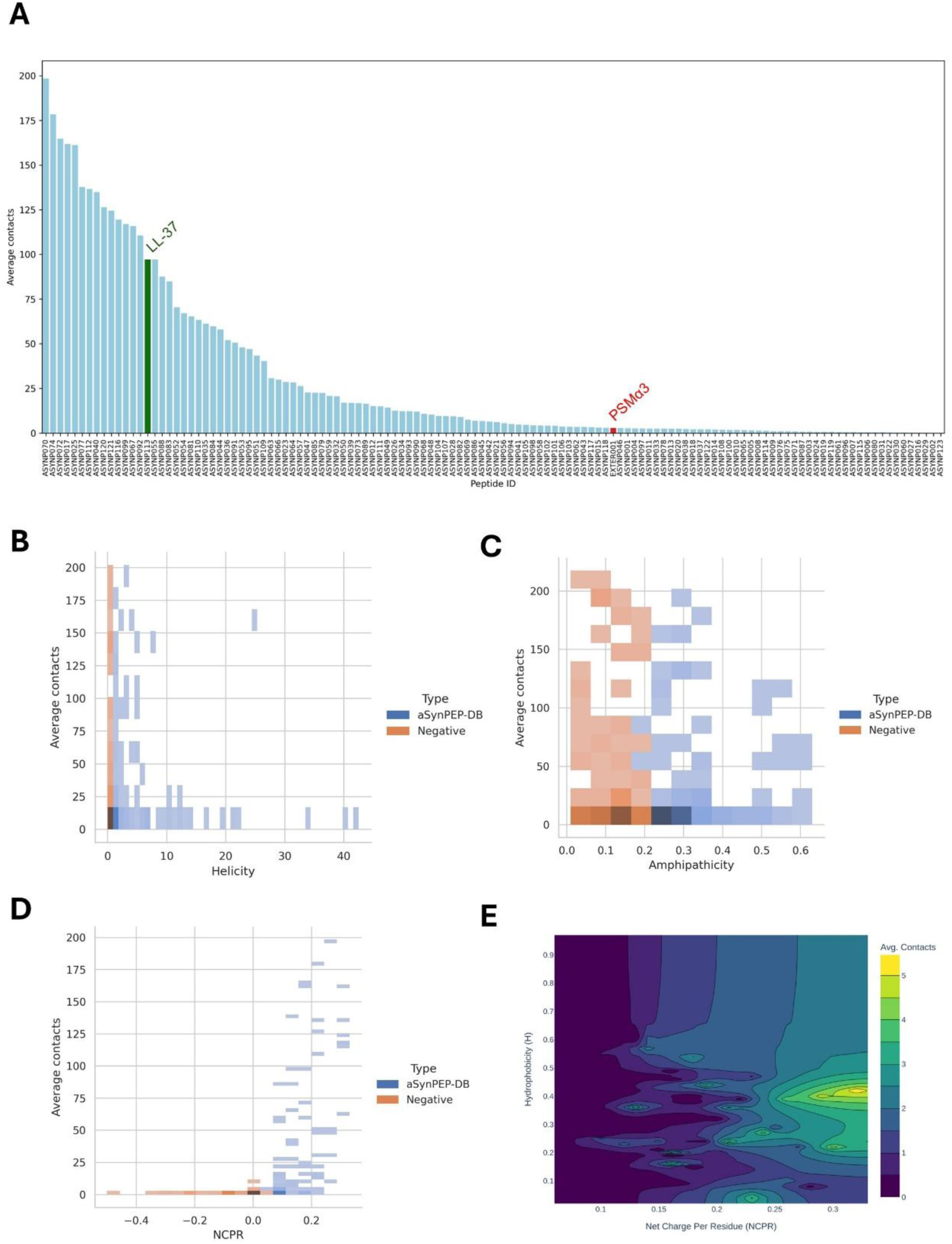
Screening and ranking of aSynPEP-DB peptides on full-length fibrils. (A) Average number of contacts between each aSynPEP-DB peptide and the full-length fibril in the “ALL” simulation; validated peptides are highlighted. (B–D) Relationships between contacts and helicity, amphipathicity, and net charge per residue (NCPR) for positives (SALL-pos from aSynPEP-DB) and negative controls (SALL-neg-hel, SALL-neg-uh and SALL-neg-ncpr). (E) Joint dependence of contacts on NCPR and hydrophobicity.

To identify which physicochemical property most strongly governs binding in this assay, we compared contact distributions for the aSynPEP-DB peptides with independent simulations of three control peptide sets, each lacking one of the defining properties used in the original curation. Neither predicted helical propensity, nor amphipathicity alone separated positives from negatives (**Fig. 8 B–C**). This result is consistent with the binding mechanism inferred above, in which peptides interact with the fibril predominantly through flexible, extended conformations. In contrast, net charge per residue (NCPR) had a dominant effect: negatively charged peptides exhibited minimal binding, whereas higher positive NCPR correlated with higher contact numbers (**Fig. 8D and SI Fig. S7**). This finding is consistent with the two-step binding mechanism resolved for the validated binders, in which electrostatic capture by the anionic fuzzy coat is a prerequisite for persistent association and hotspot engagement.

Charge, however, does not act in isolation. Although hydrophobicity alone did not correlate with binding across the positive dataset (**SI Fig. S7**), some peptides with only modest positive charge but relatively higher hydrophobicity achieved contact numbers comparable to those of more highly cationic sequences (**Fig. 8E**). This behavior indicates that optimal binding emerges from a balance between electrostatic recruitment and hydrophobic stabilization. Thus, within an already enriched candidate space, NCPR dominates the ranking, while hydrophobicity acts as a secondary modulator of binding strength.

These analyses further suggest that the predicted helical propensity and amphipathic patterning function primarily as enabling design constraints. In practice, these properties likely help maintain short hydrophobic peptides in solution and reduce their tendency to self-associate before target engagement, while preserving a residue pattern that can be deployed upon target engagement.

Consistent with this view, when LL-37 was constrained in a helical conformation (SC-helical), its contact map displayed a clearer amphipathic organization, with the hydrophobic face preferentially oriented toward the fibril core (**SI Fig. S8A**). However, despite this more ordered binding geometry, the helical peptide formed significantly fewer total contacts than the unconstrained, flexible LL-37 simulation (**SI Fig. S8B**). These results indicate that conformational adaptability, rather than a pre-formed helix, is advantageous for penetrating the fuzzy coat and maximizing multivalent interactions on the fibril surface, and suggest that the peptide becomes less helical when partitioned into the fuzzy coat. This interpretation is consistent with the observation that LL-37 remains largely disordered or only partially structured in aqueous solution (37), thereby retaining the conformational flexibility needed to adapt to fibril surfaces and sustain multivalent interactions.

## Discussion

We have constructed a model of the fibril structure of full-length aSyn and studied its conformational properties and interactions with peptides. Our results show that the surface of pathological aSyn fibrils cannot be understood simply as the exposed face of the β-sheet core. Instead, it emerges from the dynamic coupling between the ordered cross-β scaffold and the disordered fuzzy coat that surrounds it. In the brain-derived full-length fibrils simulated here, the structured core contributes recurrent aggregation-prone surfaces, whereas the disordered termini, particularly the acidic C-terminal tails, generate a highly dynamic anionic environment that transiently interact with those regions. As a consequence, the core fibril surface is neither permanently exposed nor permanently shielded but dynamically accessible through a highly charged and mobile outer layer. This view shifts the mechanistic description of aSyn fibrils from that of a rigid amyloid rod to that of a structured core embedded in a responsive polyelectrolyte brush, in which surface accessibility becomes an emergent property of the full-length assembly.

Within this framework, ligand recognition is not governed by a single pre-formed binding pocket but by a multistep, multivalent process in which long-range electrostatics and short-range hydrophobic interactions are mechanistically coupled. The fuzzy coat first acts as an electrostatic capture layer: its high density of acidic residues expands the encounter radius for cationic ligands, increases their effective local concentration near the fibril surface, and prolongs their residence time. This capture does not, in itself, generate stable binding, but enables repeated sampling of the structured core. Productive association then occurs when this confined exploration is converted into mixed electrostatic and hydrophobic stabilization on exposed aggregation-prone motifs, particularly β5/P2 and β9/P3 ladders. In this sense, the fuzzy coat functions as a funnel that converts diffuse electrostatic encounters into selective engagement of defined structural features.

This mechanism also provides a physicochemical explanation for selectivity towards aggregated aSyn. Monomeric aSyn retains its acidic character but lacks both the stacked anionic brush created by fibril assembly and the spatially organized hydrophobic ladders of the Lewy-fold core. Encounters with the monomer, therefore, remain transient, as electrostatic attraction is not reinforced by persistent capture or structured hotspots. Conversely, core-only fibrils expose hydrophobic elements but lack the electrostatic capture layer that concentrates and retains ligands at the surface. Selectivity for fibrils thus arises not from recognition of a unique folded epitope, but from recognition of an aggregate-specific mesoscale architecture that combines long-range anionic recruitment with short-range stabilization on recurrent hydrophobic surfaces.

From a therapeutic perspective, these results suggest a division of labor between physicochemical properties that govern fibril engagement. Net positive charge primarily determines electrostatic recruitment to the fibril surface, while hydrophobic residues provide the stabilization required for productive binding to core hotspots. Properties such as helical propensity or amphipathicity appear to function mainly as design constraints that maintain peptide solubility and prevent premature self-association. Consistent with this interpretation, ligands with high structural preorganization exhibit reduced binding, suggesting that flexible peptides may be better suited to exploit the heterogeneous, dynamic surface of full-length fibrils.

Although our simulations do not explicitly model fibril elongation or inhibition kinetics, the preferential localization of peptides to P2/β5 and P3/β9 is mechanistically notable. These regions correspond to the principal solvent-exposed aggregation-prone ladders of the core and have previously been implicated in fibril growth and oligomer-to-fibril transitions (36). Ligand occupancy of these surfaces, therefore, provides a plausible route by which fibril-binding peptides could interfere with amyloid formation including charge-dependent secondary nucleation (38).

More broadly, these findings support the idea that disordered flanking regions are functional components of amyloid assemblies rather than merely unresolved sequence extensions. By intermittently masking and revealing aggregation-prone motifs, the fuzzy coat is likely to influence interactions not only with ligands but also with cofactors, protein partners, and other modulators of fibril biology.

These conclusions should nevertheless be interpreted within the limits of the present framework. CALVADOS is a coarse-grained model and cannot resolve atomistic interactions such as specific hydrogen bonding, water-mediated contacts, or detailed side-chain packing. The interaction hotspots identified here should therefore be viewed as probabilistic regions that are dynamically exposed rather than fixed atomic binding pockets often found in globular proteins. In addition, the present study focuses on a single Parkinson’s disease**-**derived Lewy-fold polymorph, whereas other synucleinopathies may present distinct surface architectures.

Taken together, our results argue that full-length fibril models are not simply more complete structural representations but qualitatively different mechanistic entities. By restoring the fuzzy coat, they reveal how disorder, electrostatics, and aggregation-prone structure cooperate to define the ligand-accessible surface of pathological aSyn. This provides a mechanistic basis for peptide selectivity and suggests that effective therapeutic strategies may need to target not only the ordered amyloid core but also the dynamic disordered architecture that regulates access to it.

## Materials and Methods

### Initial structures: brain-derived aSyn fibril core and full-length reconstruction

The ordered core of aSyn filaments derived from patients with PD and dementia with Lewy bodies was obtained from the Protein Data Bank (PDB: 8A9L) (15). The deposited atomic model spans residues 31 to 100 of aSyn and represents a single-protofilament cross-β architecture. To reconstruct an extended fibril, we used the helical parameters reported with the structure (axial rise 4.76 Å and angular rotation 0.86°) to build a 180°-twist fibril. The resulting assembly contains 209 aSyn chains.

The unresolved N- and C-terminal segments were modeled to generate full-length chains. Full-length aSyn sequence information was obtained from UniProt (P37840). Disordered N- and C-terminal regions were constructed using MODELLER (39, 40) with the experimental core as a template and with positional restraints along the fibril axis (z coordinate) during loop building to minimize steric clashes between adjacent layers. The final model contains an ordered core (residues 31–100) and flexible terminal segments forming a solvent-exposed fuzzy coat.

### Peptide selection and starting conformations

We focused on two peptides with prior experimental validation as inhibitors of aSyn aggregation, PSMα3 and LL-37 (23, 24), and on a broader candidate set (n = 123) from aSynPEP-DB (25). Peptide sequences were taken from aSynPEP-DB. Initial peptide conformations were generated with ColabFold (41). Because these peptides are expected to fluctuate between disordered and partially ordered states in solution, we treated them as disordered polymers in the coarse-grained simulations, except in a dedicated control where LL-37 was restrained in an α-helical conformation.

### Simulation systems and nomenclature

Thirteen systems were prepared for simulations (**Table 1**). Two production simulations contained the full-length fibril with either PSMα3 (SP-PSM) or LL-37 (SP-LL37) at a 2:1 aSyn:peptide ratio to reflect substoichiometric inhibition conditions. Control simulations served to validate the physicochemical properties required for binding aSyn fibrils via two PSMα3 variants: all hydrophobic residues substituted by Leu (SC-all-Leu) and a charge reversion to an anionic scaffold (SC-anionic). Further controls probed the role of the fibrillar scaffold and the fuzzy coat: LL-37 with monomeric aSyn chains treated as full IDPs (SC-monomer), LL-37 with a core-only fibril lacking disordered termini (SC-core), and LL-37 restrained in a helical conformation (SC-helical). Additional controls simulated the fibril alone (SC-fibril-alone) and LL-37 alone (SC-LL37-alone). For higher-content screening, we simulated all 123 aSynPEP-DB peptides simultaneously with the fibril (SALL-pos; one copy of each peptide), and three “ALL-negative” simulations containing peptides that fail one of the defining inhibitory properties (low helicity, non-amphipathic, or negatively charged; SALL-neg-hel, SALL-neg-uh, and SALL-neg-ncpr; n = 123 peptides for each simulation) that were obtained from the list of negative peptides not meeting the screening requirements for the inclusion in aSynPEP-DB (25). Screen simulations contained one copy of each peptide to maintain a substoichiometric ratio.

### Coarse-grained molecular dynamics simulations

Simulations were performed with the CALVADOS 3 implicit-solvent coarse-grained force field (27, 29) implemented in OpenMM (42). Unless stated otherwise, simulations were carried out in the NVT ensemble at 293.15 K, pH 7.5, ionic strength 0.15 M, in a cubic box of side length 120 nm. Simulations were run for 1 μs using a Langevin integrator with a time step of 10 fs and a friction coefficient of 0.01 ps^−1^. Configurations were saved every 100 ps. GPU acceleration was enabled.

To preserve the experimentally determined cross-β core while allowing conformational sampling of the fuzzy coat, we restrained residues 31 to 100 in each aSyn chain through an elastic network: harmonic potentials with force constant 700 kJ mol^−1^ nm^−2^ are applied between residues whose centers of mass are separated by less than 0.9 nm in the initial structure. N- and C-terminal segments were unrestrained. The charges of N- and C-terminal residues of aSyn and peptides were adjusted to account for the positively charged amino group and the negatively charged carboxylate group. In the core-only fibril control, the truncated aSyn segments were left uncharged at both ends to reflect a cleaved section and avoid artificial end charges.

### Contact analysis

Trajectory analyses were performed with MDTraj (43) v1.10.1. The first 100 ns were discarded as equilibration (**SI Fig. S9**). For each frame, the center of mass (COM) of each aSyn chain and each peptide was calculated. Residue contacts were computed for aSyn–peptide chain pairs with COM distances below 15 nm. Contacts were scored with a continuous hyperbolic tangent switching function that yields a contact value of 0.5 at an inter-residue distance of 1.0 nm (**SI Fig. S10**). Contact maps were obtained by averaging across frames and peptides.

To classify binding events for PSMα3 and LL-37, we used a threshold of 40 simultaneous contacts to label frames as “significantly interacting” (**SI Fig. S11**). For intrafibril aSyn contacts, the terminal four chains at each end of the fibril were excluded to minimize finite-size end effects. For each aSyn chain, contacts with neighboring chains up to four positions above and below were computed and summed with self-contacts. Statistical significance between datasets was assessed with Mann-Whitney-Wilcoxon two-sided test. Electrostatic interactions were defined as contacts between acidic residues (Asp, Glu) and basic residues (Arg, Lys). Hydrophobic interactions were defined as contacts among Val, Leu, Ile, Met, Phe, Trp, and Pro.

### Solvent-accessible surface area

Per-residue SASA was computed with MDTraj (43) using the Shrake and Rupley algorithm (44). To focus on a representative bulk region, the top 10 and bottom 10 chains were removed prior to analysis. Radii were adjusted to reflect the coarse-grained representation using a water radii of 0.14 nm and residue sigma values for CALVADOS amino acid representation from (45). The standard error of the mean was computed from block averaging across the trajectory.

### Aggregation propensity and aggregation hotspots

Sequence-based aggregation propensity was predicted with AGGRESCAN (32). Regions with at least five consecutive residues with positive aggregation propensity values were considered aggregation hotspots. Structural aggregation propensity was predicted with Aggrescan4D (34) on ten core chains extracted from the fibril center, using stability calculations and a distance of aggregation of 10 Å. Both the top and bottom chains were removed to avoid finite-size end effects.

### Peptide binding orientation

To quantify the preferred orientation of peptides on the fibril surface, we computed the principal axis of each peptide using principal component analysis. The eigenvector associated with the largest eigenvalue of the covariance matrix of residue coordinates was taken as the peptide direction vector, and its angle relative to the fibril axis was used as a measure of binding orientation.

## Data availability

Simulation trajectories and topologies of all systems are available at https://doi.org/10.34810/data2621. Data and scripts to reproduce the analyses are available at https://github.com/PPMC-lab/aSyn-fibrils-MD.

## Acknowledgments

CP-G was supported by the European Molecular Biology Organization (EMBO_SEG11214), the Secretariat of Universities and Research of the Catalan Government (2023 FI_3 00018), and by the UAB (2025PILIFRUA37). OB was supported by the Spanish Ministry of Science and Innovation via a doctoral grant (FPU22/03656). SV was supported by the Spanish Ministry of Science and Innovation (PID2022-137963OB-I00), ICREA, ICREA-Academia 2020 and 2021-SGR-00635 AGAUR (Generalitat de Catalunya) and CERCA Programme (Generalitat de Catalunya). We also acknowledge support from the Novo Nordisk Foundation challenge program PRISM (Protein Interactions and Stability in Medicine and Genomics, NNF18OC0033950, to K.L.-L.)

## References

1. E. R. Dorsey, T. Sherer, M. S. Okun, B. R. Bloem, The Emerging Evidence of the Parkinson Pandemic. J Parkinsons Dis 8, S3–S8 (2018).

2. G. N. D. C. Group, Global, regional, and national burden of neurological disorders during 1990-2015: a systematic analysis for the Global Burden of Disease Study 2015. Lancet Neurol 16, 877–897 (2017).

3. T. Pardo-Moreno et al., Current Treatments and New, Tentative Therapies for Parkinson’s Disease. Pharmaceutics 15 (2023).

4. I. Alafuzoff, P. Hartikainen, Alpha-synucleinopathies. Handb Clin Neurol 145, 339–353 (2017).

5. M. Goedert, R. Jakes, M. G. Spillantini, The Synucleinopathies: Twenty Years On. J Parkinsons Dis 7, S51–S69 (2017).

6. M. G. Spillantini et al., Alpha-synuclein in Lewy bodies. Nature 388, 839–840 (1997).

7. F. X. Theillet et al., Structural disorder of monomeric α-synuclein persists in mammalian cells. Nature 530, 45–50 (2016).

8. T. S. Ulmer, A. Bax, N. B. Cole, R. L. Nussbaum, Structure and dynamics of micelle-bound human alpha-synuclein. J Biol Chem 280, 9595–9603 (2005).

9. N. Cremades et al., Direct observation of the interconversion of normal and toxic forms of α-synuclein. Cell 149, 1048–1059 (2012).

10. A. J. Dear et al., Kinetic diversity of amyloid oligomers. Proc Natl Acad Sci U S A 117, 12087–12094 (2020).

11. R. Gaspar et al., Secondary nucleation of monomers on fibril surface dominates α-synuclein aggregation and provides autocatalytic amyloid amplification. Q Rev Biophys 50, e6 (2017).

12. T. R. Jahn et al., The common architecture of cross-beta amyloid. J Mol Biol 395, 717–727 (2010).

13. A. W. Fitzpatrick et al., Atomic structure and hierarchical assembly of a cross-β amyloid fibril. Proc Natl Acad Sci U S A 110, 5468–5473 (2013).

14. M. Schweighauser et al., Structures of α-synuclein filaments from multiple system atrophy. Nature 585, 464–469 (2020).

15. Y. Yang et al., Structures of α-synuclein filaments from human brains with Lewy pathology. Nature 610, 791–795 (2022).

16. S. H. W. Scheres, B. Ryskeldi-Falcon, M. Goedert, Molecular pathology of neurodegenerative diseases by cryo-EM of amyloids. Nature 621, 701–710 (2023).

17. S. Ansari, et al., In cell NMR reveals cells selectively amplify and structurally remodel amyloid fibrils. bioRxiv (2024).

18. M. Milanesi, Z. F. Brotzakis, M. Vendruscolo, Transient interactions between the fuzzy coat and the cross-β core of brain-derived Aβ42 filaments. Sci Adv 11, eadr7008 (2025).

19. M. Cullen et al., Integrating NMR Restraints into Coarse-Grained Simulations: Toward Accurate Conformational Ensembles of Complex Protein Systems. J Am Chem Soc (2026).

20. S. Pacheco et al., Phosphorylation of α-Synuclein Fibrils at S129 Changes DNAJB1 Binding as Probed by Solid-State NMR. JACS Au 6, 343–356 (2026).

21. Y. Han et al., Fibril fuzzy coat is important for α-synuclein pathological transmission activity. Neuron 113, 1723–1740.e1727 (2025).

22. V. Iglesias, O. Bárcenas, C. Pintado-Grima, M. Burdukiewicz, S. Ventura, Structural information in therapeutic peptides: Emerging applications in biomedicine. FEBS Open Bio (2024).

23. J. Santos et al., α-Helical peptidic scaffolds to target α-synuclein toxic species with nanomolar affinity. Nat Commun 12, 3752 (2021).

24. J. Santos, I. Pallarès, S. Ventura, Is a cure for Parkinson’s disease hiding inside us? Trends Biochem Sci 47, 641–644 (2022).

25. C. Pintado-Grima et al., aSynPEP-DB: a database of biogenic peptides for inhibiting α-synuclein aggregation. Database (Oxford) 2023 (2023).

26. C. Pintado-Grima, S. Ventura, The role of amphipathic and cationic helical peptides in Parkinson’s disease. Protein Sci 34, e70020 (2025).

27. G. Tesei, T. K. Schulze, R. Crehuet, K. Lindorff-Larsen, Accurate model of liquid-liquid phase behavior of intrinsically disordered proteins from optimization of single-chain properties. Proc Natl Acad Sci U S A 118 (2021).

28. G. Tesei, K. Lindorff-Larsen, Improved predictions of phase behaviour of intrinsically disordered proteins by tuning the interaction range. Open Res Eur 2, 94 (2022).

29. F. Cao, S. von Bülow, G. Tesei, K. Lindorff-Larsen, A coarse-grained model for disordered and multi-domain proteins. Protein Sci 33, e5172 (2024).

30. S. von Bülow, K. E. Johansson, K. Lindorff-Larsen, AF-CALVADOS: AlphaFold-guided simulations of multi-domain proteins at the proteome level. bioRxiv, 2025.2010.2019.683306 (2025).

31. P. Tompa, Structural disorder in amyloid fibrils: its implication in dynamic interactions of proteins. FEBS J 276, 5406–5415 (2009).

32. O. Conchillo-Solé et al., AGGRESCAN: a server for the prediction and evaluation of “hot spots” of aggregation in polypeptides. BMC Bioinformatics 8, 65 (2007).

33. S. Errico et al., Structural commonalities determined by physicochemical principles in the complex polymorphism of the amyloid state of proteins. Biochem J 482, 87–101 (2025).

34. O. Bárcenas et al., Aggrescan4D: structure-informed analysis of pH-dependent protein aggregation. Nucleic Acids Res 52, W170–W175 (2024).

35. R. Sim et al., Fluorescence Detection of Alpha-Synuclein Aggregates in the Gut Using a Peptide Probe. ACS Chem Neurosci 17, 1211–1226 (2026).

36. J. Santos et al., A Targetable N-Terminal Motif Orchestrates α-Synuclein Oligomer-to-Fibril Conversion. J Am Chem Soc 146, 12702–12711 (2024).

37. J. Johansson, G. H. Gudmundsson, M. E. Rottenberg, K. D. Berndt, B. Agerberth, Conformation-dependent antibacterial activity of the naturally occurring human peptide LL-37. J Biol Chem 273, 3718–3724 (1998).

38. A. K. Buell et al., Solution conditions determine the relative importance of nucleation and growth processes in α-synuclein aggregation. Proc Natl Acad Sci U S A 111, 7671–7676 (2014).

39. B. Webb, A. Sali, Comparative Protein Structure Modeling Using MODELLER. Curr Protoc Bioinformatics 54, 5.6.1–5.6.37 (2016).

40. A. Sali, T. L. Blundell, Comparative protein modelling by satisfaction of spatial restraints. J Mol Biol 234, 779–815 (1993).

41. M. Mirdita et al., ColabFold: making protein folding accessible to all. Nat Methods 19, 679–682 (2022).

42. P. Eastman et al., OpenMM 8: Molecular Dynamics Simulation with Machine Learning Potentials. J Phys Chem B 128, 109–116 (2024).

43. R. T. McGibbon et al., MDTraj: A Modern Open Library for the Analysis of Molecular Dynamics Trajectories. Biophys J 109, 1528–1532 (2015).

44. A. Shrake, J. A. Rupley, Environment and exposure to solvent of protein atoms. Lysozyme and insulin. J Mol Biol 79, 351–371 (1973).

45. Y. C. Kim, G. Hummer, Coarse-grained models for simulations of multiprotein complexes: application to ubiquitin binding. J Mol Biol 375, 1416–1433 (2008).

