## Supporting Information for "Full-Length Molecular Models of Brain-Derived α-Synuclein Fibrils Reveal a Fuzzy-Coat-Mediated Mechanism for Selective Peptide Binding"

**Supplementary Figures**

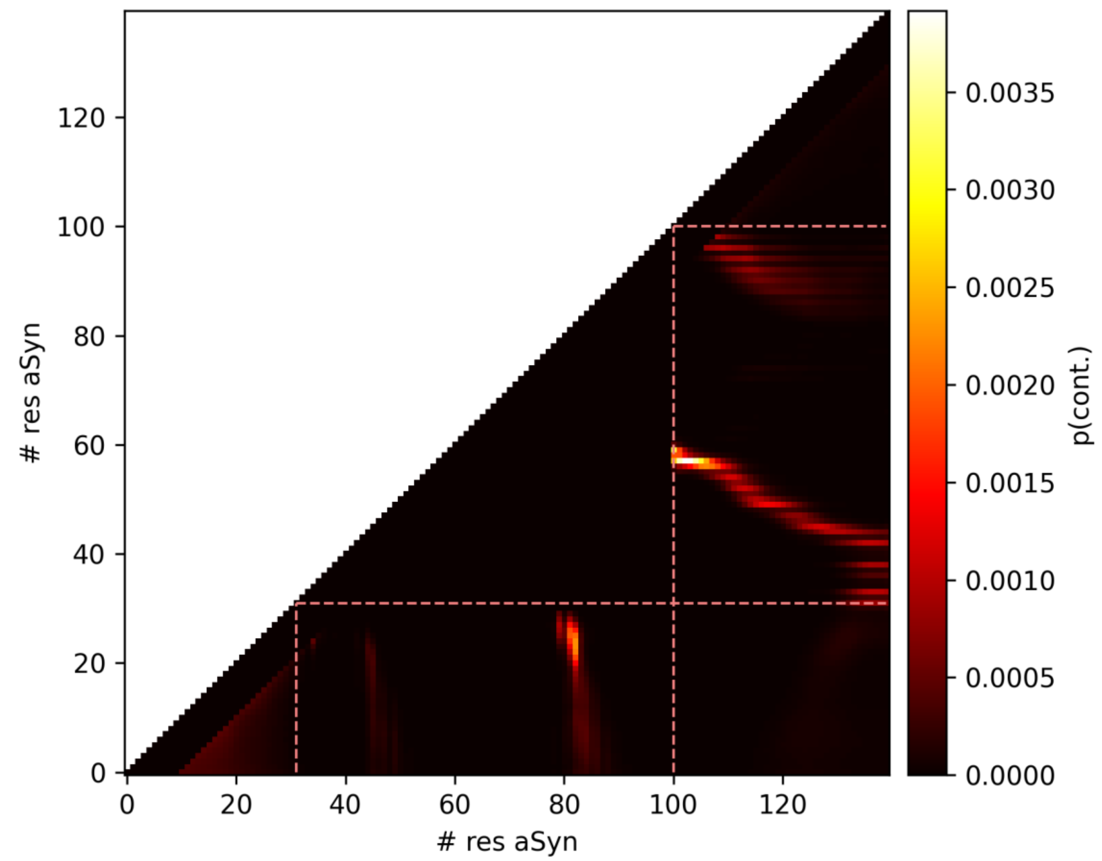


**Fig. S1.** aSyn self contacts between IDRs and the core. Contacts are observed for both N- and C-termini. N-t IDR interacts with the hinge region between β8 and β9 whereas C-t contacts overlap with β5 and β9 core regions.

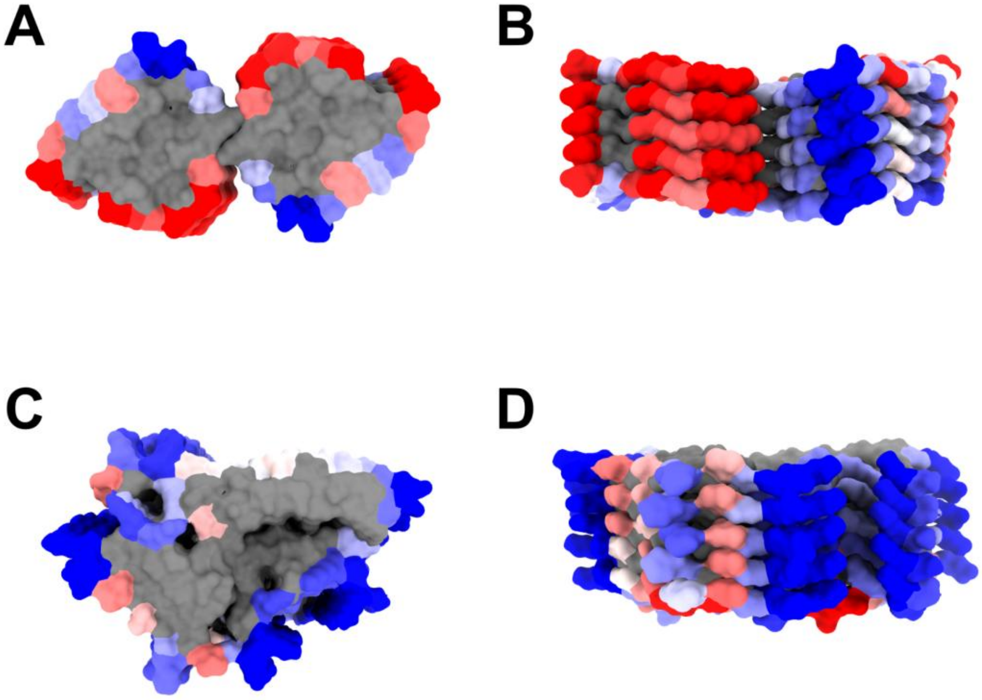


**Fig. S2.** A4D comparison between five middle chains of Aβ42 fibrils type II (PDB ID: 7Q4M) and aSyn fibrils (PDB ID: 8A9L). (A) Top view of Aβ42. (B) Lateral view of Aβ42. (C) Top view of aSyn. (D) Lateral view of aSyn. Red, blue, white, and grey colors indicate aggregation-prone, soluble, neutral, and no solvent exposition, respectively.

A

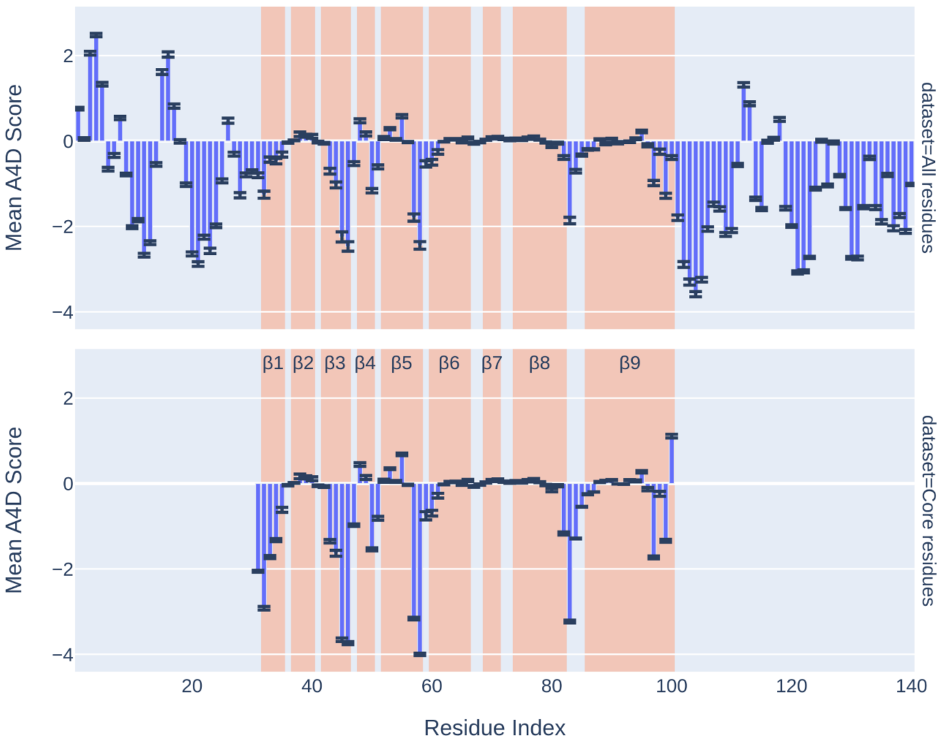


B

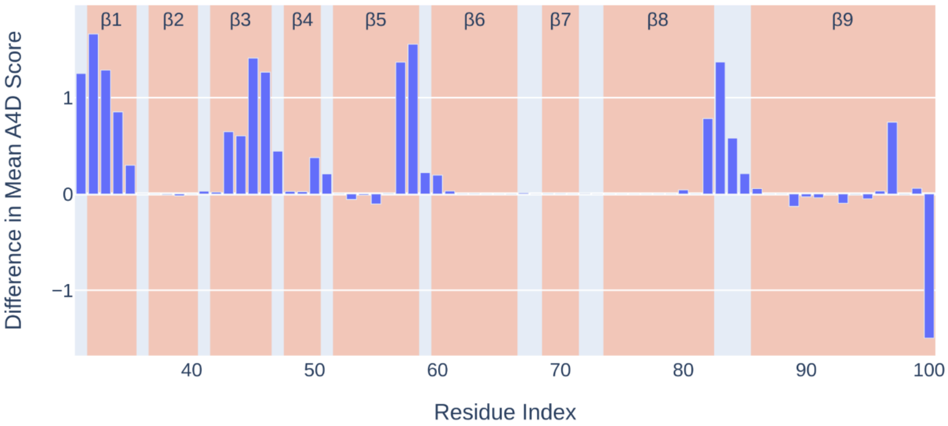


**Fig. S3.** Per-residue A4D scores in aSyn fibril. (A) Mean A4D score for the full-length aSyn fibril and for core-only residues. Standard Error of the Mean shown as error bars. (B) Difference in mean A4D score for core residues in the presence and absence of the fuzzy coat. The light-red background indicates the location of β-strands.

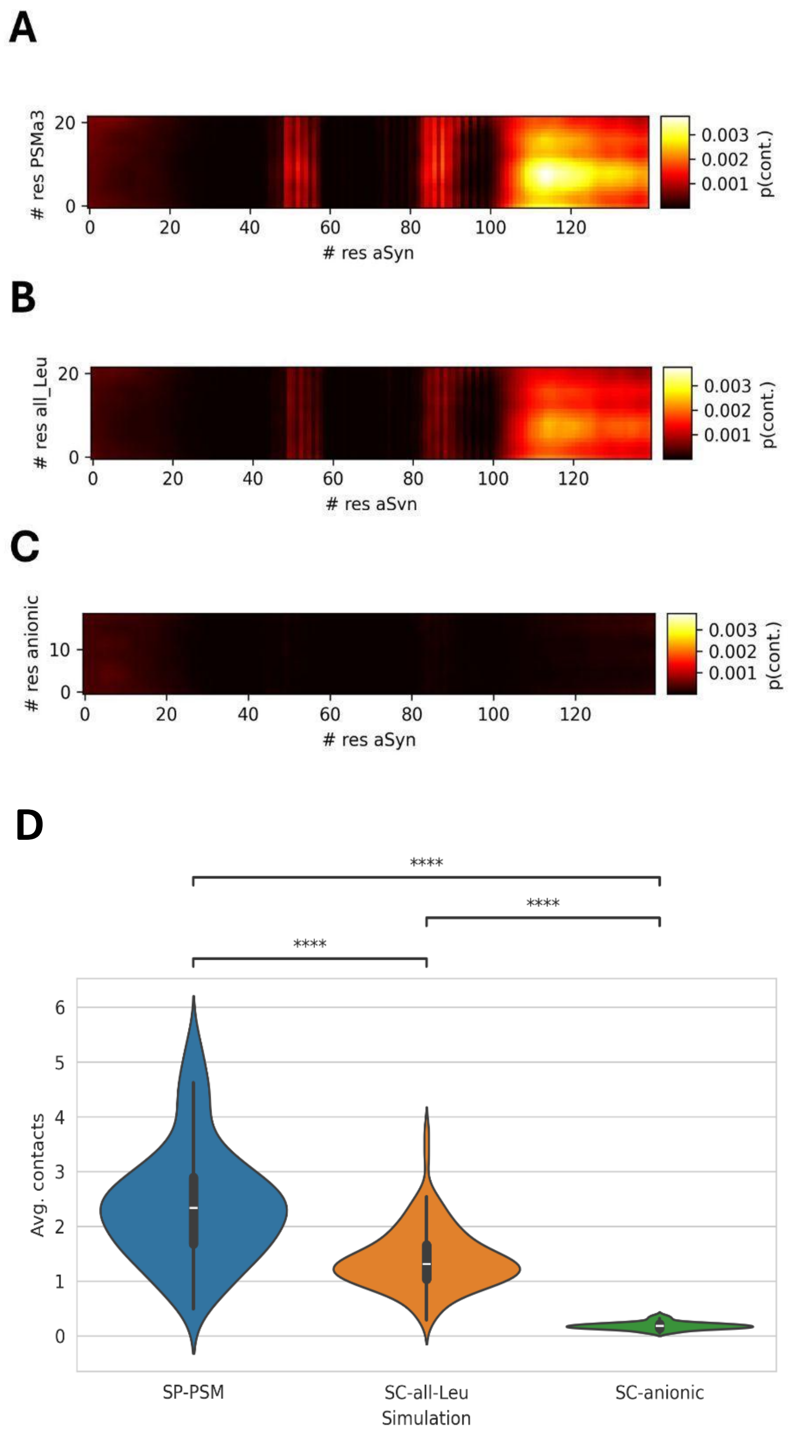


**Fig. S4.** PSMa3 variants contacts. (A) Contact map of original PSMa3 peptide with aSyn fibrils. (B) All-Leu variant of PSMa3 displays similar contacts than PSMa3, engaging with both C-terminal tail and inner core regions of aSyn fibrils. (C) The anionic variant does interact with aSyn fibrils given the electrostatic repulsion with the fuzzy coat. (D) Difference in average contacts between production simulation with original PSMa3 peptide and the two variant controls; nominally, **** indicates p<2.014e-14 for a difference in average contacts. The simulated systems are described in Table 1.

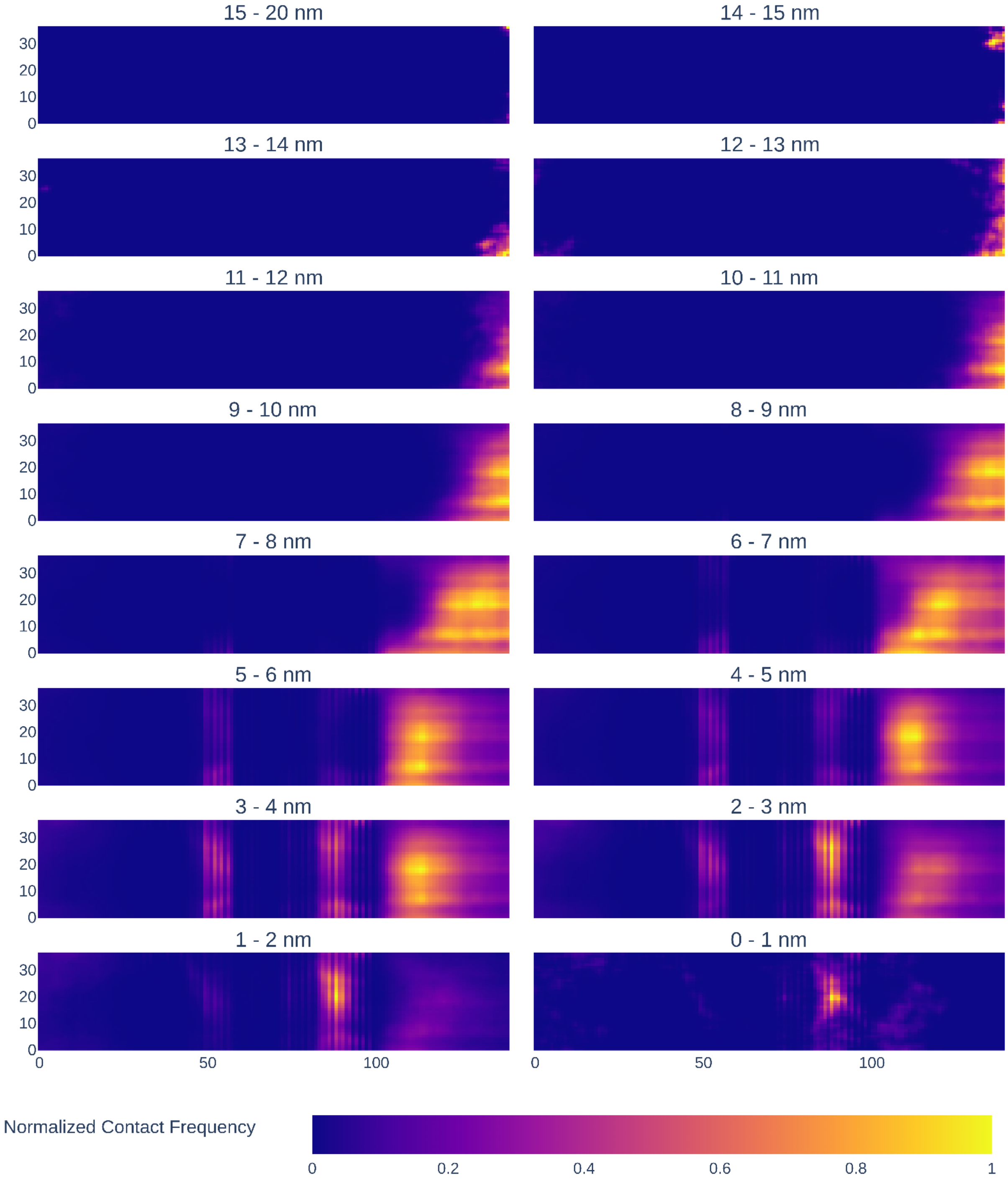


**Fig. S5.** LL37-aSyn contact maps at different peptide-aSyn COM distances from 15 nm to 0 nm in intervals of 1 nm. At intermediate distances (5-10nm) peptide interactions engage with aSyn C-terminal IDRs that approach them towards the cross-beta core of the fibril, allowing for inner contacts with the core below 5 nm.

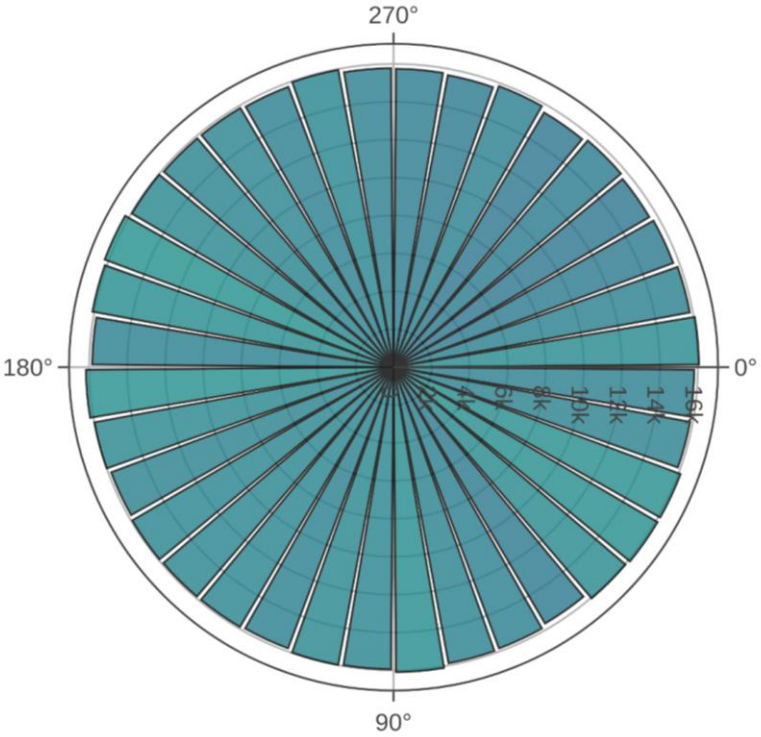


**Fig. S6**. Axis angles distribution for LL-37 peptides at distances above 15 nm from aSyn COM. Instead of the preferential perpendicular binding of LL-37 peptides with the fibrillar core when they are close to the fibril, peptides that do not interact with aSyn show random angle distributions.

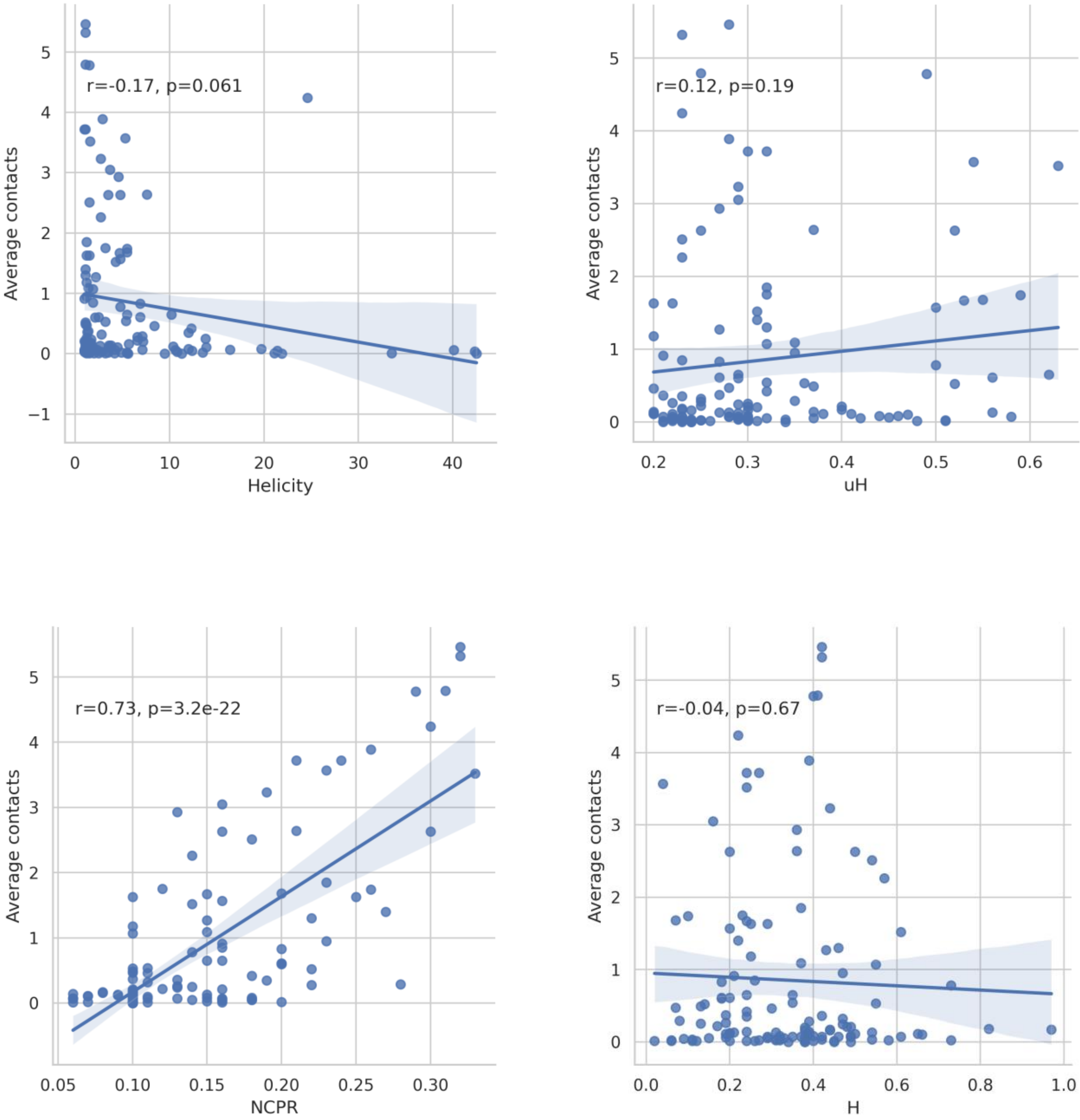


**Fig. S7.** Linear regression plots for aSynPEP-DB peptides correlating helicity, amphipathicity (uH), net charge per residue (NCPR) and hydrophobicity (H) values with average contacts. Only NCPR shows a significant correlation with a r value = 0.73.

**A**

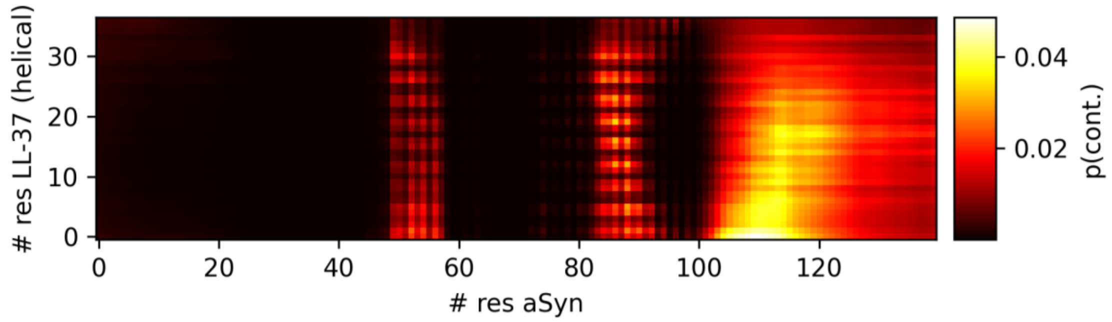


**B**

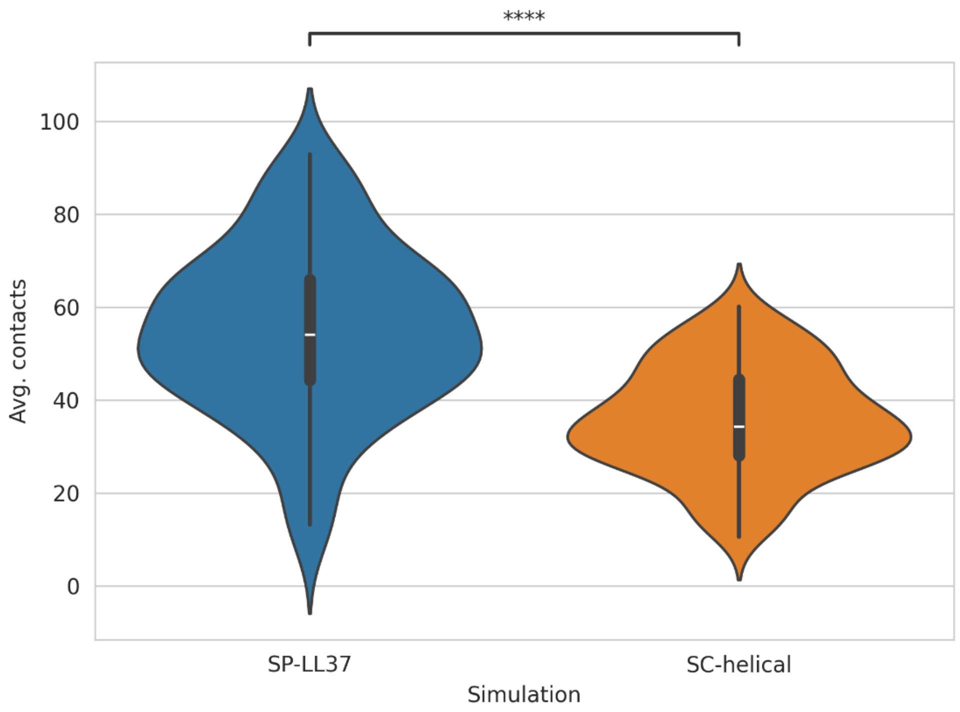


**Fig. S8**. Contact comparison between helical and disordered LL-37 peptides. (A) Contact map for LL-37 peptide with restricted secondary structure and full-length aSyn fibrils. (B) Average contact difference between disordered (SP-LL37) and helical (SC-helical) LL-37 peptides; nominally, **** indicates p=2.631e-15 for a difference in average contacts. The simulated systems are described in Table 1.

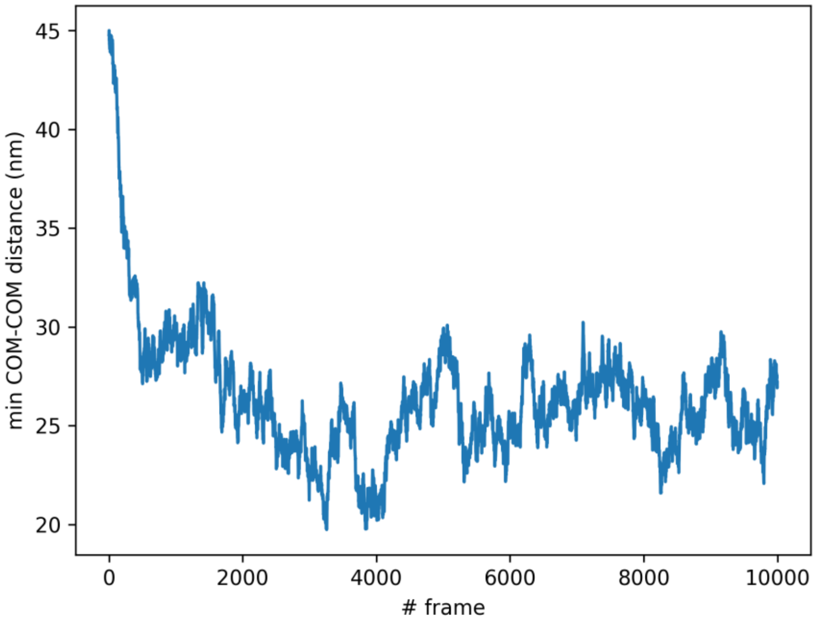


**Fig. S9.** Average minimum COM-COM distance between LL-37 peptides and aSyn chains in the fibril. After the initial 1000 frames ~100 ns, different LL-37 peptides are contacting the fibril and the system. Albeit with small fluctuations, the system could be considered equilibrated for further analyses.

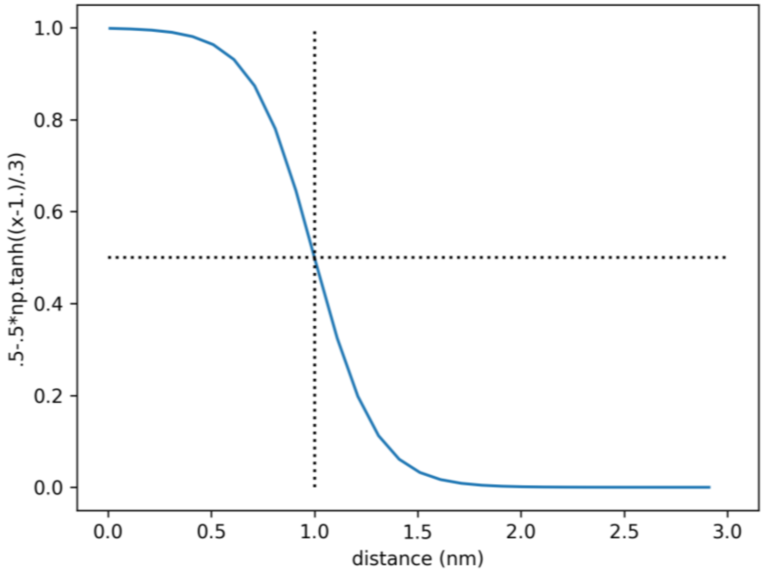


**Fig. S10.** Hyperbolic tangent function to calculate residue contacts. Two residues that are 1nm apart scores 0.5. Shorter distances rapidly score up to 1 whereas longer distances decrease to 0.

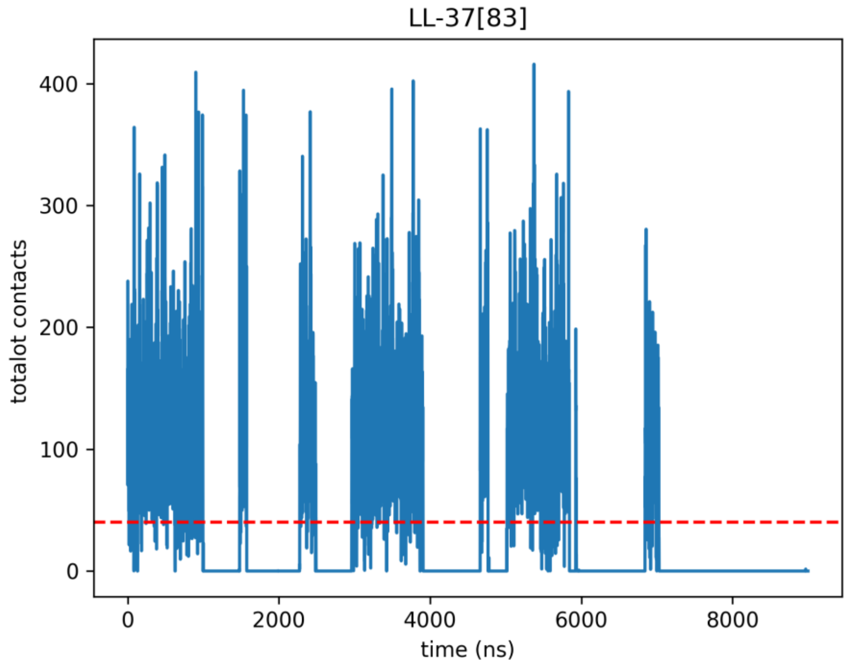


**Fig. S11.** Total number of contacts over the trajectory with aSyn fibrils for a selected LL-37 peptide. Based on the interactions, we set a discriminatory threshold at 40 (red dotted line) in order to discretize whether peptides interact or not with aSyn.

### Supplementary Tables

**Table S1.** Aggrescan4D scores of the amyloid fibril model. Both untrimmed ("With fuzzy coat") and trimmed versions ("Without fuzzy coat") were analyzed. For the full-length version, including the fuzzy coat, we computed the average Aggrescan4D scores for the complete amyloid ("Total average score") and the core region ("Core average score"); for the trimmed version, there is no difference between the complete and only-core models. The Aggrescan4D score also includes the standard error of the mean (SEM) in parentheses.

| | Total average score (SEM) | Core average score (SEM) |
|---|---:|---:|
| With fuzzy coat | -0.706 (0.006) | -0.311 (0.007) |
| Without fuzzy coat | | -0.536 (0.007) |
